# Impact of Pathogenic Variants of the Ras-MAPK Pathway on Major White Matter Tracts in the Human Brain

**DOI:** 10.1101/2024.04.17.590011

**Authors:** Monica Siqueiros-Sanchez, Erpeng Dai, Chloe A. McGhee, Jennifer A. McNab, Mira M. Raman, Tamar Green

## Abstract

Noonan syndrome (NS) and Neurofibromatosis type-1 (NF1) are genetic conditions linked to pathogenic variants in genes of the Ras-MAPK signaling pathway. Both conditions hyperactivate signaling of the Ras pathway and exhibit a high prevalence of neuropsychiatric disorders. Further, animal models of NS and NF1 and human imaging studies show white-matter abnormalities in both conditions. While these findings suggest Ras pathway hyper-activation effects on white-matter, it is unknown whether these effects are syndrome-specific or pathway-specific. To characterize the effect of NS and NF1 on human white-matter microstructural integrity and discern potential syndrome-specific influences on microstructural integrity of individual tracts, we collected diffusion-weighted imaging data from children with NS (n=24), NF1 (n=28), and age and sex-matched controls (n=31). We contrasted the clinical groups (NS or NF1) and controls using voxel-wise, tract-based, and along-tract analyses. Outcomes included voxel-wise, tract-based and along-tract fractional anisotropy (FA), axial diffusivity (AD), radial diffusivity (RD) and mean diffusivity (MD). NS and NF1 showed similar patterns of reduced FA and increased AD, RD, and MD on white-matter relative to controls and different spatial patterns. NS presented a more extensive spatial effect than NF1 on white-matter integrity as measured by FA. Tract-based analysis also demonstrated differences in effect magnitude with overall lower FA in NS compared to NF1 (d=0.4). At the tract-level, NS specific-effects on FA were detected in association tracts (superior longitudinal, uncinate and arcuate fasciculi; p’s <0.012) and NF1 specific-effects were detected in the corpus callosum (p’s<0.037) compared to controls. Results from along-tract analyses aligned with results from tract-based analyses and indicated that effects are pervasive along the affected tracts. In conclusion, we find that pathogenic variants in the Ras-MAPK pathway are associated with white-matter abnormalities as measured by diffusion in the developing brain. Overall NS and NF1 show common effects on FA and diffusion scalars, as well as specific unique effects, namely on temporoparietal-frontal tracts (intra-hemispheric) in NS and on the corpus callosum (inter-hemispheric) in NF1. The observed specific effects not only confirm prior observations from independent cohorts of NS and NF1 but also inform on syndrome-specific susceptibility of individual tracts. Thus, these findings suggest potential targets for precise, brain-focused outcome measures for existing medications, such as MEK inhibitors, that act on the Ras pathway.

## Introduction

The Ras–extracellular signal-regulated kinase mitogen-activated protein kinase signaling cascade (Ras-MAPK pathway) is crucial for brain development.^1–3^ Germline mutations in this pathway result in a collection of disorders with partially overlapping clinical features known as Rasopathies.^3^ Noonan syndrome (NS) and Neurofibromatosis type-1 (NF1) are the most common, affecting 1:2000 and 1:3000 individuals, respectively.^4,5^ NS is caused by mutations in genes encoding upregulating components of the Ras-MAPK pathway, e.g., *PTPN11, SOS1, RAF1, RIT1, BRAF, KRAS, HRAS, MEK1/2, SOS2,* and *LZTR1*.^3,6^ NF1 is caused by mutations in *NF1,* the gene encoding the neurofibromin protein and a negative regulator of the Ras-MAPK pathway.^7^ Preclinical studies in mice show that causal mutations of NS and NF1 upregulate Ras signaling and its downstream pathways leading to abnormal myelination in the brain.^8,9^ Gain-of-function mutations of protein Shp2 (encoded by *PTPN11*), an upstream effector of Ras, and of ERK1/2, a downstream effector of Ras, result in fewer myelinated axons, myelinating glial cell progenitors and mature oligodendrocytes, and axonal degeneration.^10^ Loss-of-function mutations in NF1 result in decompaction of the myelin sheath^11^ and increased numbers of myelinating glial progenitor cells.^12^

Neuroimaging diffusion voxel-wise studies show that, relative to typically developing individuals (TD), individuals with NS present widespread lower fractional anisotropy (FA) and higher axial diffusivity (AD) and radial diffusivity (RD),^13^ whereas individuals with NF1 present widespread lower FA,^14^ higher RD^15^ and mean diffusivity (MD),^16^ as well as lower FA in the corpus callosum.^17^ These findings suggest shared effects of Ras-MAPK pathway activation on white-matter. However, these studies have only independently contrasted NS and NF1 to TD on white-matter microstructure but not to each other. Therefore, whether NS and NF1 present syndrome-specific effects is unknown.

Research on white-matter microstructure in tracts beyond the corpus callosum is limited in both syndromes. Therefore, whether certain tracts are more susceptible to NS-associated genes (*PTPN11, SOS1) or* to the effects of *NF1* is unknown. For example, how NS and NF1 affect white-matter integrity association tracts (tracts connecting brain regions within the same hemisphere) is limited. Association tracts are key for social processes including mentalizing and empathy^18,19^ and deficits in social processes are highly prevalent in NS and NF1.^20,21^ Furthermore, the extent of the effect of NS and NF1 along these tracts is unclear. That is, whether white-matter microstructure is compromised along the entirety of the tract or localized at specific segments.

In this study, we utilize a rigorous design to directly compare the effects on structural brain connectivity of two distinct populations, NS and NF1, in the developing brain. Our analysis contrasts prepubertal children with confirmed pathogenic variants in the *PTPN11* or *SOS1* genes (associated with NS) and the *NF1* gene (associated with NF1) against age- and sex-matched controls. We aim to determine: (1) whether NS and NF1 exhibit common or distinct effects on FA and diffusion scalars (AD, RD, and MD), (2) whether NS and NF1 exhibit common or distinct effects on specific white-matter tracts, and (3) determine the specific locations along the association tracts where differences are evident.

## Methods

### Participants

Recruited participants (age 6-13 years) included fifteen children with NS, sixteen children with NF1, and fifteen TD controls. Individuals with NS or NF1 provided proof of genetic testing demonstrating the presence of mutations in *PTPN11* (*n*=20), *SOS1* (*n*=2), or in *NF1*. Pubertal status was ascertained using Tanner staging.^22,23^ Exclusion criteria included premature birth (<34 weeks), birth weight <2000g, history of head trauma with loss of consciousness or neurological disorders known to affect cognitive development or brain structure (e.g., seizures), known presence of gliomas in cerebellum, brainstem, or basal ganglia, and magnetic resonance imaging (MRI) contraindications. To maximize study compliance participants required a full-scale intelligence quotient (IQ) ≧70. NS and NF1 are typically associated with low-to-average IQ, therefore a cut-off of ≧70 is unlikely to exclude a significant portion of these populations. After assessing the quality of the imaging data (NS= 24; NF1= 28; TD= 31), sixty-seven individuals were included in this study (NS=22, NF1=18, TD=26).

Participants were recruited from May 2021 to January 2023. NS and NF1 participants were recruited nationwide via the Stanford University School of Medicine’s website, the Children’s Tumor Foundation, the Noonan Syndrome Foundation, Rasopathies Network and Rasopathies social media groups, relevant physician referrals and social media advertising. TD participants were recruited from local networks and social media advertising restricted to the Bay area and its surrounding areas. Legal guardians provided written informed consent for all participants. Participants aged >7 provided a complementary written assent. The authors assert that all procedures contributing to this work comply with the ethical standards of the relevant national and institutional committees on human experimentation and with the Helsinki Declaration of 1975, as revised in 2008. All procedures involving human subjects/patients were approved by the Stanford University School of Medicine Institutional Review Board.

### Behavioral assessments

Cognitive ability was ascertained using the *<u>Wechsler Abbreviated Scale of Intelligence</u>* <u>|</u> *2^nd^ Edition*^24^; we assessed overall intelligence quotients, performance and verbal intelligence quotients. To confirm that social deficits were also present in the current sample, we assessed social processes using the *Social Responsiveness Scale 2^nd^ Edition* (SRS-2).^25^ The SRS-2 is a valid 65-item parental rating scale assessing a continuum of social impairment severity, including subthreshold variation levels of autistic-like traits. Higher scores in the SRS-2 indicate higher symptom severity.

### Image acquisition and preprocessing

All participants completed behavioral training in a mock MRI scanner prior undergoing the real MRI scan as described in prior studies^26,27^ and in the supplementary. MRI scanning was completed on a GE Premier 3.0 Tesla whole-body system using a standard 48-channel head coil (GE Healthcare, Milwaukee, WI).

Diffusion-weighted images were collected using a single-shot diffusion-weighted spin-echo echo-planar multi-shell imaging sequence with a series of b=0 scans in the encoding and reversed phase encoding polarity for bulk and distortion correction. Preprocessing of diffusion-weighted images corrected susceptibility-induced and eddy current distortions and motion artifacts with FSL’s topup^28^ and eddy.^29–32^ Whole-brain high-resolution T1-weighted magnetization-prepared rapid gradient-echo (MPRAGE) images were collected during the same session. During each acquisition, researchers monitored subject motion and assessed scan quality. Scans were repeated if needed. T1 weighted images were preprocessed with the standard Freesurfer 7.2.0 recon-all pipeline (https://surfer.nmr.mgh.harvard.edu). Technical details of this pipeline are described in prior publications.^33–37^ Independent raters (MS, MR, CM) examined the corrected diffusion weighted images and the Freesurfer-generated gray/white and pial surfaces. Pial surfaces were manually edited when necessary and re-generated after edits. Imaging parameters and further details on acquisition and quality control of imaging data can be found in the Supplementary.

### Image analysis

Diffusion imaging metrics (i.e., FA, AD, RD, MD) capture the random motion of water molecules in biological tissues and provide insights into white-matter microstructural properties including tissue organization, integrity, and connectivity. FA measures the degree of directionality of water diffusion within a voxel and ranges from 0-1 (values closer to zero indicate a low degree of diffusion asymmetry). FA is thought to capture different underlying biological factors of white-matter, including degree of coherence of axonal bundles, axon density, diameter, and myelination.^38^ However, since FA is a summary measure, examining other diffusion metrics provides additional information. AD measures water diffusivity parallel to the main diffusion direction. RD measures water diffusivity orthogonal to the main diffusion direction. MD is the average of the apparent diffusion coefficient of all directions. There are no direct correlates between diffusion scalars and biological causes of disease, however higher AD is linked to higher axonal integrity^39^ and lower RD to higher axonal myelination.^40^

### Tract-Based Spatial Statistics (TBSS)

Voxel-wise analysis of whole-brain FA, AD, RD, and MD data were carried out using the TBSS^41^ standard pipeline in FSL.^42^ TBSS projects all subjects’ FA, AD, RD, and MD data onto a mean FA tract skeleton that represents the center of all tracts common to all subjects before applying voxel-wise cross-subject statistics. Further details on TBSS can be found in the Supplementary.

### TRActs Constrained by UnderLying Anatomy (TRACULA)

TRACULA automatically delineates 42 major white-matter tracts using probabilistic tractography.^43^ We extracted weighted-means of FA, RD, AD and MD from each of the 42 tracts after TRACULA’s 3D-tract-reconstruction. Weighted means are means computed from diffusion scalars in each voxel weighted by the probability of said voxel being part of a specific tract.

Tracts that failed to adequately reconstruct or did so partially were excluded (Table S1).^44^ The groups did not differ on head motion measures (see supplementary).^45^ Further details of the TRACULA toolbox and quality control procedures can be found in the Supplementary.

### Pointwise Assessment of Streamline Tractography Attributes (PASTA)

As singular values for each tract, tract-based measures from TRACULA (weighted means) may “average out” important but fine-grained between-group differences located along different tract parts. To identify where these putative differences were located along each tract, within-group averaged point values of FA, AD, RD, and MD were generated from consecutive cross-sections of each tract using PASTA (included in TRACULA) and compared between the groups. Tracts included in these analyses per group can be found in Table S1.

### Statistical analysis

We performed whole-brain voxel-wise analyses of skeletonized FA, MD, AD, and RD data across groups (NS vs. TD; NF1 vs. TD; NS vs. NF1) using TBSS and FSL *randomize*^46^ (5000 permutations). We adjusted for multiple comparisons and family-wise error rates using Threshold-Free Cluster Enhancement (significance threshold p<.05), considering age and sex as covariates.

Tract-based analyses focused on the corpus callosum, anterior thalamic radiation, frontal aslant tract, arcuate, inferior longitudinal, middle longitudinal, superior longitudinal (I, II, III), and uncinate fasciculi. We selected these tracts based on TBSS results (areas of potential convergence and divergence between clinical groups), evidence from previous studies (i.e., corpus callosum in NF1), and relevance for higher order processes in Rasopathies.^13^ We examined between-group differences in FA, AD, RD, and MD with cohen’s *d* effect sizes and analysis of covariance with age and sex as covariates and correcting for multiple comparisons using the false discovery rate (FDR). Follow-up analyses contrasted clinical groups to TD and to each other using t-tests.

Fine-grained between-group differences in FA, AD, RD, and MD of consecutive cross-sections along each tract were examined using general linear models embedded in PASTA, with age and sex as covariates and adjusting for multiple comparisons using FDR. Statistical analyses outside pipeline-embedded tools were performed in RStudio 4.0.5 (RStudio, PBC); analytic code supporting the findings is available from the corresponding author upon reasonable request.

## Results

### Demographic and behavioral

Table 1 presents this sample’s demographic, cognitive, social responsiveness data, pubertal stage, and medication information. The groups did not differ on age, sex, or pubertal stage. Both clinical groups had lower FSIQ (p’s<.005) on the Wechsler Abbreviated Scale of Intelligence 2^nd^ Ed. We also confirmed social deficits in our sample as clinical groups exhibited higher scores on the SRS-2 (p’s < .002) than the TD group; clinical groups did not differ on IQ nor SRS-2 scores (Table 1).

**Table 1.**
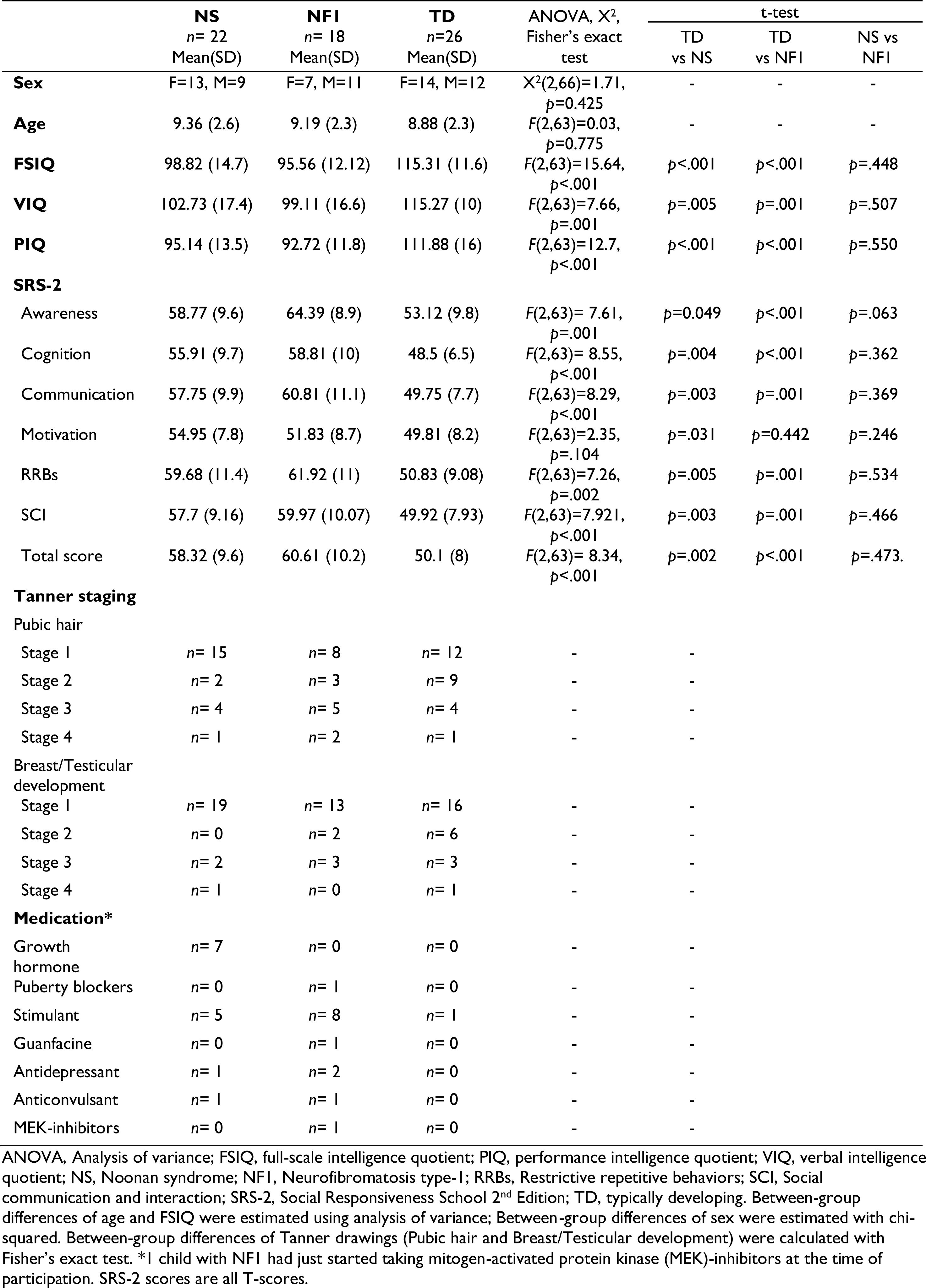
Descriptive groupwise statistics of included participants.

|  | **NS**<br>n= 22<br>Mean(SD) | **NFI**<br>n= 18<br>Mean(SD) | **TD**<br>n=26<br>Mean(SD) | ANOVA, X<sup>2</sup>,<br>Fisher's exact<br>test | t-test |  |  |
| --- | --- | --- | --- | --- | --- | --- | --- |
|  |  |  |  |  | TD<br>vs NS | TD<br>vs NFI | NS vs<br>NFI |
| **Sex** | F=13, M=9 | F=7, M=11 | F=14, M=12 | X<sup>2</sup>(2,66)=1.71,<br>p=0.425 | - | - | - |
| **Age** | 9.36 (2.6) | 9.19 (2.3) | 8.88 (2.3) | F(2,63)=0.03,<br>p=0.775 | - | - | - |
| **FSIQ** | 98.82 (14.7) | 95.56 (12.12) | 115.31 (11.6) | F(2,63)=15.64,<br>p<.001 | p<.001 | p<.001 | p=.448 |
| **VIQ** | 102.73 (17.4) | 99.11 (16.6) | 115.27 (10) | F(2,63)=7.66,<br>p=.001 | p=.005 | p=.001 | p=.507 |
| **PIQ** | 95.14 (13.5) | 92.72 (11.8) | 111.88 (16) | F(2,63)=12.7,<br>p<.001 | p<.001 | p<.001 | p=.550 |
| **SRS-2** |  |  |  |  |  |  |  |
| Awareness | 58.77 (9.6) | 64.39 (8.9) | 53.12 (9.8) | F(2,63)= 7.61,<br>p=.001 | p=0.049 | p<.001 | p=.063 |
| Cognition | 55.91 (9.7) | 58.81 (10) | 48.5 (6.5) | F(2,63)= 8.55,<br>p<.001 | p=.004 | p<.001 | p=.362 |
| Communication | 57.75 (9.9) | 60.81 (11.1) | 49.75 (7.7) | F(2,63)=8.29,<br>p<.001 | p=.003 | p=.001 | p=.369 |
| Motivation | 54.95 (7.8) | 51.83 (8.7) | 49.81 (8.2) | F(2,63)=2.35,<br>p=.104 | p=.031 | p=0.442 | p=.246 |
| RRBs | 59.68 (11.4) | 61.92 (11) | 50.83 (9.08) | F(2,63)=7.26,<br>p=.002 | p=.005 | p=.001 | p=.534 |
| SCI | 57.7 (9.16) | 59.97 (10.07) | 49.92 (7.93) | F(2,63)=7.921,<br>p<.001 | p=.003 | p=.001 | p=.466 |
| Total score | 58.32 (9.6) | 60.61 (10.2) | 50.1 (8) | F(2,63)= 8.34,<br>p<.001 | p=.002 | p<.001 | p=.473 |
| **Tanner staging** |  |  |  |  |  |  |  |
| Pubic hair |  |  |  |  |  |  |  |
| Stage 1 | n= 15 | n= 8 | n= 12 | - | - |  |  |
| Stage 2 | n= 2 | n= 3 | n= 9 | - | - |  |  |
| Stage 3 | n= 4 | n= 5 | n= 4 | - | - |  |  |
| Stage 4 | n= 1 | n= 2 | n= 1 | - | - |  |  |
| Breast/Testicular<br>development |  |  |  |  |  |  |  |
| Stage 1 | n= 19 | n= 13 | n= 16 | - | - |  |  |
| Stage 2 | n= 0 | n= 2 | n= 6 | - | - |  |  |
| Stage 3 | n= 2 | n= 3 | n= 3 | - | - |  |  |
| Stage 4 | n= 1 | n= 0 | n= 1 | - | - |  |  |
| **Medication\*** |  |  |  |  |  |  |  |
| Growth<br>hormone | n= 7 | n= 0 | n= 0 | - | - |  |  |
| Puberty blockers | n= 0 | n= 1 | n= 0 | - | - |  |  |
| Stimulant | n= 5 | n= 8 | n= 1 | - | - |  |  |
| Guanfacine | n= 0 | n= 1 | n= 0 | - | - |  |  |
| Antidepressant | n= 1 | n= 2 | n= 0 | - | - |  |  |
| Anticonvulsant | n= 1 | n= 1 | n= 0 | - | - |  |  |
| MEK-inhibitors | n= 0 | n= 1 | n= 0 | - | - |  |  |

ANOVA, Analysis of variance; FSIQ, full-scale intelligence quotient; PIQ, performance intelligence quotient; VIQ, verbal intelligence quotient; NS, Noonan syndrome; NFI, Neurofibromatosis type-1; RRBs, Restrictive repetitive behaviors; SCI, Social communication and interaction; SRS-2, Social Responsiveness School 2<sup>nd</sup> Edition; TD, typically developing. Between-group differences of age and FSIQ were estimated using analysis of variance; Between-group differences of sex were estimated with chi-squared. Between-group differences of Tanner drawings (Pubic hair and Breast/Testicular development) were calculated with Fisher's exact test. \*1 child with NFI had just started taking mitogen-activated protein kinase (MEK)-inhibitors at the time of participation. SRS-2 scores are all T-scores.

### NS and NF1 affect white-matter integrity

Voxel-wise analyses showed widespread differences between clinical groups and TD individuals on all diffusion scalars across the corpus callosum, anterior thalamic radiation, and association tracts. NS and NF1 groups showed lower global FA in comparison to the TD group (NS *p=*.0002; NF1 *p=*.0002) (Fig. 1). Visual inspection of FA images suggested NS affected FA more pervasively than NF1 across the white matter skeleton. In addition, certain tracts were primarily affected by NS (e.g., superior longitudinal fasciculi) and others by NF1 (e.g., the corpus callosum).

**Figure 1.**
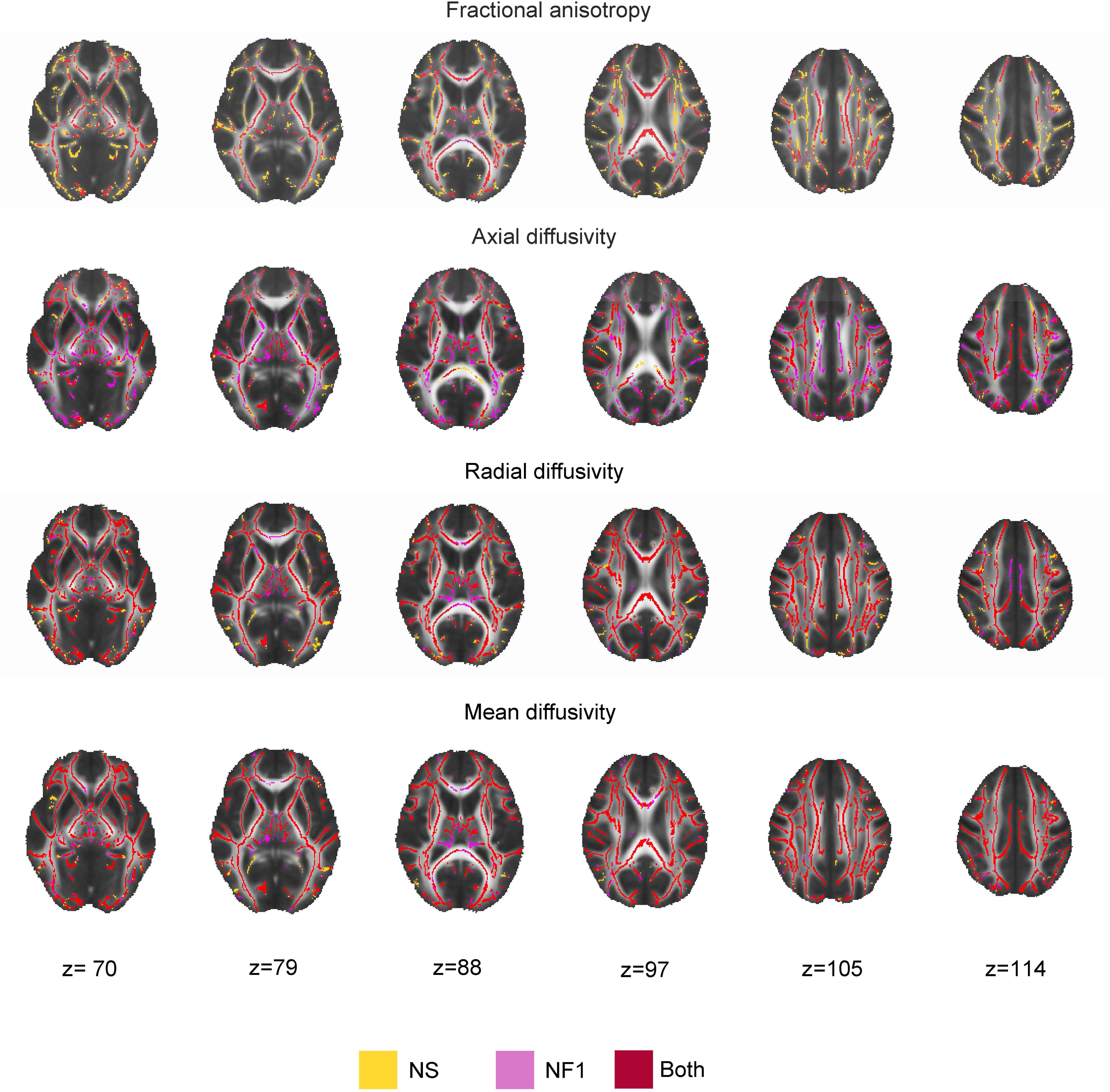
Noonan syndrome (NS) and Neurofibromatosis type-1 (NF1) have pervasive effects on whole-brain white matter. Whole-brain voxel-wise differences (*p<*.05, FDR-adjusted) between clinical groups and typically developing (TD) individuals while controlling for age and sex. Overall, clinical groups showed lower fractional anisotropy (FA) and higher diffusivities (axial diffusivity[AD]; radial diffusivity[RD]; mean diffusivity[MD]) than TD individuals. Voxels where FA, AD, RD or MD are significantly different between only one clinical group and TD group are colored in magenta (NF1) or yellow (NS); voxels where both clinical groups are significantly different from TD controls are colored in red. (A) Lower FA in clinical groups relative to controls. (B) Higher AD in clinical groups relative to controls. (C) Higher RD in clinical groups relative to controls. (D) Higher MD in clinical groups relative to controls.

Individuals with NS or NF1 had higher AD, RD, and MD than the TD group (NS *p’s<*.001; NF1 *p’s<*.001; FDR-corrected) (Fig. 1). Visual inspection of voxel-wise images suggested differences in spatial effects of NS and NF1 on long association tracts. For example, the FA in the superior longitudinal fasciculus appeared more pervasively affected by NS than by NF1 with corresponding more perversive effect on AD in the NF1 compared to NS (Fig. 1). RD and MD images suggested a similar spatial effects of both syndromes on most white-matter tracts in these scalars (Fig. 1).

FDR-corrected statistical analyses showed individuals with NS presented lower global FA (*p=*0.0495) than individuals with NF1, whereas individuals with NF1 had higher global AD (*p=* 0.0008), MD (*p=*0.0294) and higher RD (*p=*0.047) than NS.

### Spatial and magnitude-dependent impact of NS and NF1 on distinct white-matter tracts

Next, we conducted tract-based analyses of white-matter scalars in the corpus callosum, anterior thalamic radiation, and association tracts (Fig. 1E). Table S2 presents scalar means, standard deviations, and cohen’s *d* effect sizes. Tract-based analyses in FA, AD, RD, and MD showed a main effect of group on most of the tracts (Table S3).

When comparing the clinical groups with the TD group, we found similar patterns across the clinical groups: a decrease in FA and an increase in AD, RD, and MD. These shared patterns were observed in the corpus callosum (FA: central, temporal, and parietal body, genu, and splenium; AD: parietal, premotor, and temporal body), right anterior thalamic radiation, bilateral frontal aslant tract, and most ventral association tracts (inferior and middle longitudinal fasciculi) (Fig.2; Table S3). RD and MD were higher in NS and NF1 than in the TD group on most tracts (Table S3; Fig.S1).

**Figure 2.**
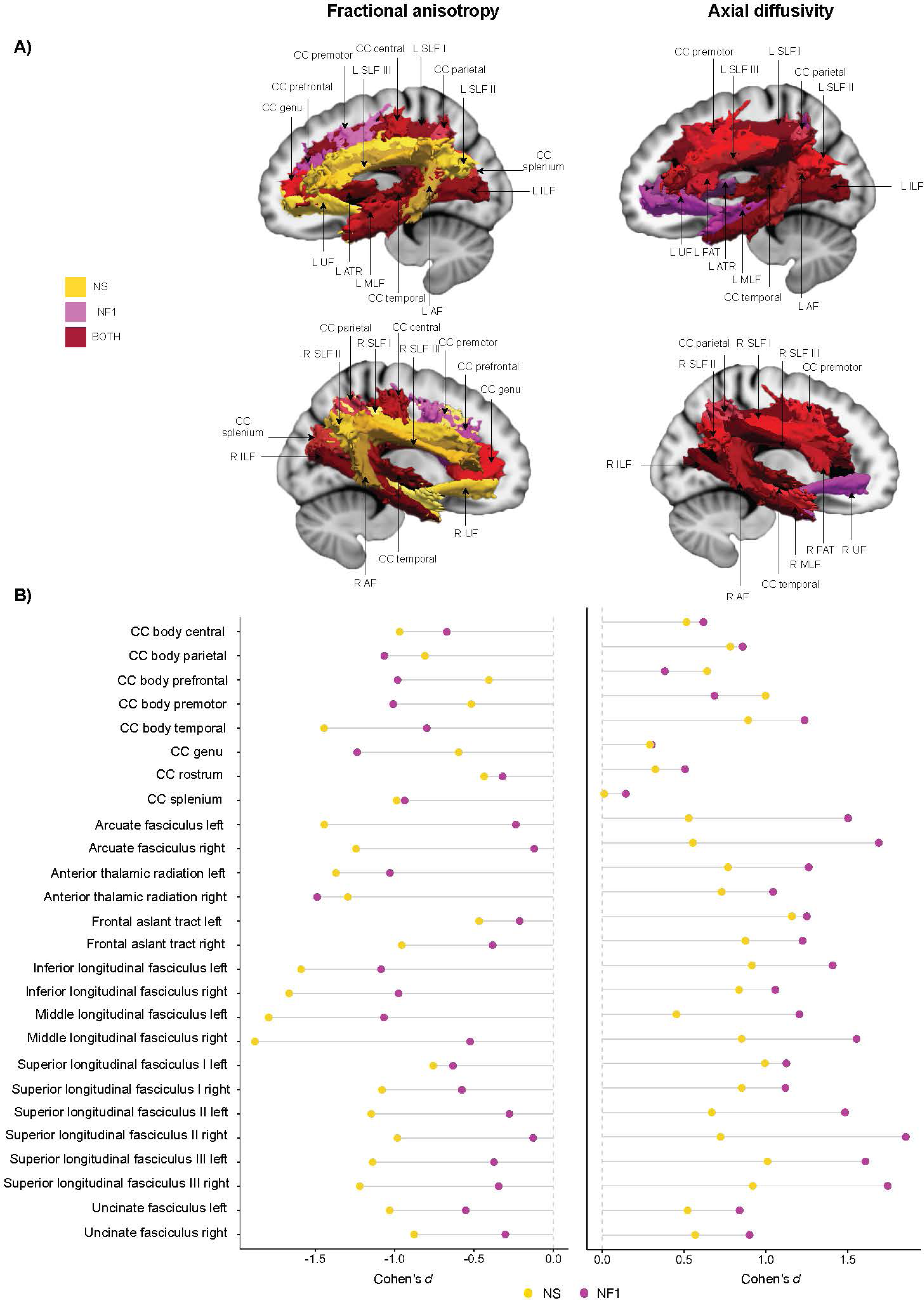
Noonan syndrome (NS) primarily affects FA on long association tracts whereas Neurofibromatosis type-1 (NF1) primarily affects the corpus callosum and AD. Tract based effects of NS and NF1 on fractional anisotropy (FA) and axial diffusivity (AD) on commissural, projection, and association tracts. (A) Sections of the brain’s corpus callosum and connecting tracts are color-coded to show where FA (left) or AD (right) is impacted in clinical groups. Yellow indicates tracts affected by NS, magenta for NF1, and red for tracts impacted by both NS and NF1 in any metric (FA, AD, RD, or MD) (B) Effect sizes (cohen’s *d*) of mean differences between clinical groups and TD in FA (left) and AD (right) from select commissural, projection, and association tracts. NS’s cohen’s *d* values are depicted as yellow circles and NF1’s cohen’s *d* values are depicted as magenta circles. Analysis of covariance and t-tests results can be found in table S3. FA and AD means, standard deviations, effect sizes and 95% confidence intervals can be found in table S2. AF: Arcuate Fasciculus; ATR: Anterior Thalamic Radiation; CC: Corpus Callosum; FAT: Frontal Aslant Tract; ILF: Inferior Longitudinal Fasciculus; MLF: Middle Longitudinal Fasciculus; NF1: Neurofibromatosis type-1; NS: Noonan syndrome; R: Right; SLF: Superior Longitudinal Fasciculus; UF: Uncinate Fasciculus.

In comparison to typically developing (TD) individuals, distinct effects specific to each syndrome emerged, characterized by unique spatial and magnitude patterns. In the NS group, FA was primarily reduced in the uncinate, arcuate, and superior longitudinal fasciculi. This reduction in FA was not evident in the NF1 group (see Figure 2A and Table S3). Conversely, the NF1 group showed a significant impact on AD within the uncinate fasciculus, implying more intact structural connectivity in these regions compared to the NS group (see Figures 2.A-B and Table S3). The scale of these spatial differences was confirmed by the effect sizes (Table S2).

Regarding the magnitude of the changes, the effect sizes illustrated in Figure 2.B and Table S2 indicate that NS had a consistently greater impact on FA in the long association tracts (both ventral and superior) than NF1, when compared to controls. Conversely, within the corpus callosum, FA was more significantly affected by NF1 than by NS, relative to controls.

The homogeneity of variance was not met for the RDs and MDs of several tracts. Therefore, we conducted a sensitivity analysis examining tract-based between-group differences in white matter using non-parametric Kruskal-Wallis and Mann-Whitney.^47^ Results from non-parametric analyses aligned with the analysis of covariance results, affirming the validity of our findings (data not shown).

### Along-tract between-group differences in association tracts

Tract-based analyses may “average out” differences at specific locations of a tract. Thus, follow-up exploratory analyses examined between-group differences on FA and AD along each association tract with PASTA. We centered our study on FA and AD measurements specifically within association tracts. This decision was based on the observed differences in spatial patterns and matrices patterns between the clinical groups.

Overall, our along-tract analyses supported the findings from our tract-based analysis. Specifically, for association tracts tested FA was primary affected in NS (Fig. 3A). Meanwhile, AD changes were prominent in the uncinate fasciculus for the NF1 group as described but also in larger parts of the longitudinal fasciculus. Specifically, compared to TD, AD was affected in large sections of the superior longitudinal fasciculus III in NF1 (Fig. 3B).

**Figure 3.**
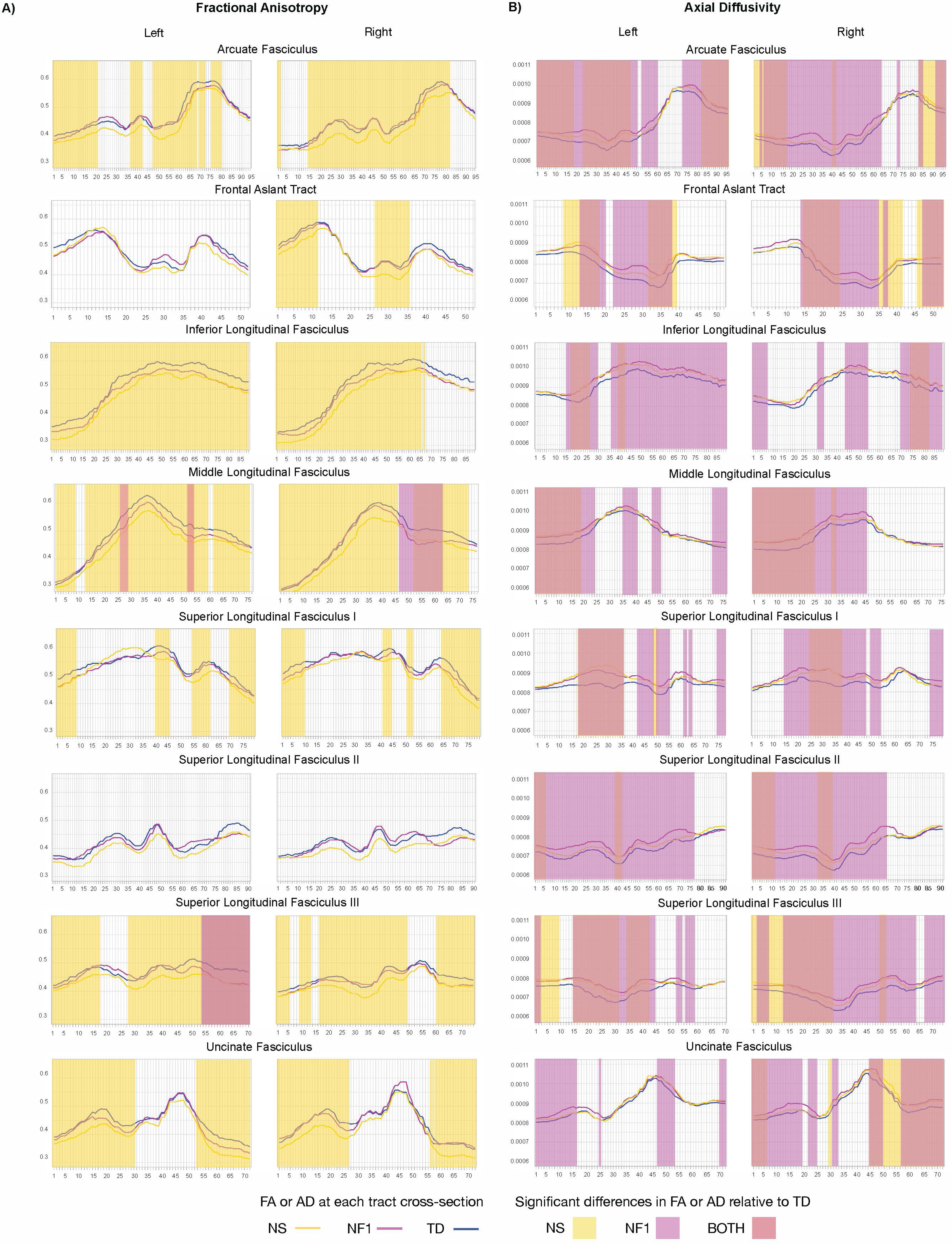
Along-tract locations of reduced mean fractional anisotropy (FA) and increased mean axial diffusivity (AD) in NS and NF1 of select association tracts. Follow-up analyses assessing fine-grained between-group differences in FA and AD of consecutive cross-sections along select association tracts using general linear models, considering age and sex as covariates, embedded in Pointwise Assessment of Streamline Tractography Attributes (PASTA) framework. (A) Mean between-group differences in FA along association tracts. (B) Mean between-group differences in AD along association tracts. Mean FA or AD values of consecutive cross-section of each tract are depicted as colored lines: the NS group is displayed in yellow, the NF1 group is displayed in magenta, and the TD group in blue. Cross-sections where there are significant differences (FDR-adjusted) in mean FA (A) or mean AD (B) between a clinical group and the TD group are signaled by rectangles: differences between NS and TD are depicted in yellow, differences between NF1 and TD are depicted in magenta, differences between both clinical groups (NS and NF1) and the TD group overlap are displayed in red. FA or AD values are depicted along the *y* axis and are metric specific, ranging between 0.3-0.65 for FA and 0.0006-0.0011 for AD; values along the *x* axis are tract specific and dependent on the length of the tract as they reflect the number of cross-sections sampled from each tract. AD: axial diffusivity; FA: fractional anisotropy. NF1: Neurofibromatosis type-1; NS: Noonan syndrome; TD: typically developing.

Certain between-group differences from tract-based averages of FA and AD did not replicate in these analyses. We did not detect differences in FA scalar in the superior longitudinal fasciculus I (for NF1, FA averages were lower than controls) and II (for NS, FA averages were lower than controls)(Fig. 3A). Despite these discrepancies, results from along-tract analyses overall mirrored those from tract-based analyses, showing that significant parts of the tract were compromised and confirming the pervasive effect of NS on FA and NF1 on AD.

## Discussion

This study examined the effects of pathogenic variants in genes associated with Ras-MAPK pathway on white-matter microstructure in the human developing brain. Voxel-wise analyses showed pervasive reduced FA in children with NS and NF1 compared to TD, indicating compromised structural connectivity. Overall, NS and NF1 displayed overlapping spatial patterns; yet specifically, NS impacted FA more broadly than NF1. Complementary tract-based analysis further showed distinct spatial patterns of differences and effect magnitude in FA of each syndrome: compared to controls, FA was reduced in NS, but not in NF1 in intra-hemispheric long association tracts (superior longitudinal, uncinate and arcuate fasciculus). In contrast, FA was reduced in inter-hemispheric tracts in parts of the corpus callosum in NF1 but not in NS. NS and NF1 groups exhibited higher AD, RD, and MD relative to the TD group across most tracts (voxel-wise and tract-based), suggesting compromised white matter microstructure integrity in both syndromes. Lastly, along-tract evaluations of association tracts mirrored the patterns seen in tract-based analyses: FA disparities were more pronounced in NS compared to NF1. The increased AD in NF1 relative to NS within these association tracts implies relatively preserved structural connectivity along these tracts’ main direction in NF1, leading to a less compromised FA in NF1 than in NS. The observed specific effects not only confirm prior observations from independent cohorts of NS and NF1 but also suggest potential targets for precise, brain-focused, outcome measures for existing medications that act on the Ras pathway such as MEK inhibitors. Finally, these findings of impaired structural connectivity add to the emerging evidence implicating the Ras-MAPK pathway in human white-matter integrity as well as in higher order cognitive processes subserved by these tracts.

Our findings suggest compromised myelin in both NS and NF1 as evidenced by consistently lower FA values compared to controls. Compromised myelin is likely due to impaired myelin formation. Mice models show evidence of abnormalities in oligodendrocyte lineal cells resulting from mutations in *PTPN11* and *NF1,*^10,48,49^ suggesting that compromised myelin could underpin the white-matter alterations we observe in NS and NF1. Specifically, gain-of-function mutations in *Ptpn11* result in hypomyelination, fewer myelinated axons and abnormalities of the myelin sheath^10^ which may explain the lower FA we observed in NS. In NF1, *Nf1* mutant mice show abnormal myelin wrapping in axons within the brain.^50^ Similarly, neurofibromin is highly expressed in oligodendrocytes, and research in NF1 shows abnormalities in oligodendrocyte lineal cells, including aberrant specification and proliferation of oligodendrocyte-precursor cells^48,49^ that can affect myelination and reduce FA. In addition to FA, we detected increased AD, RD, and MD in most brain tracts, which suggests widespread alterations. These alterations may be linked to the Ras-MAPK pathway and further support the hypothesis of myelin damage in children with Rasopathies. AD reflects water movement along nerve fibers, RD across fibers which may signal myelin loss, and MD represents overall water diffusion within the brain tissue.^38,40,51^ Our observations align with prior research that has documented similar white matter changes in specific brain regions for NS and NF1. Notably, in NS, key brain structures showed reduced FA and increased RD.^13^ Further, in NF1 increased RD and MD in the corpus callosum is a consistent finding in humans^15–17^ and in *Nf1* mutant mice.^50,52^ An additional study observed increased AD, RD, and MD in the superior longitudinal fasciculus in individuals with NF1.^53^ These consistent patterns across studies underscore the potential impact of Ras-MAPK pathway overactivation on white-matter integrity.

In addition to these shared effects of NS and NF1 on white matter integrity across various tracts, our analysis identified that the syndromes differ in their spatial influence. Specifically, NS predominantly affects long association tracts, while NF1 has a more extensive impact on the corpus callosum. In terms of magnitude, effects on FA were pronounced in NS and modest in NF1 across almost all tracts, except in certain areas of the corpus callosum where the impact of NF1 on FA exceeds that of NS. AD was higher in NF1 relative to NS across all association tracts except for the corpus callosum where NF1 exhibited lower FA, relative to NS. NF1 mice models show decompaction of the myelin sheath in axons in the corpus callosum^50,52^ which may explain the reduced FA and the profound effects observed in this white-matter bundle. These findings suggest a stronger impact of neurofibromin in inter-hemispheric tracts. The intra-hemispheric tracts, the arcuate, superior longitudinal, and uncinate fasciculi are implicated in action production, emotional intelligence, social cognition (i.e. mentalizing), and language in the general population.^18,19,62,63^ In NF1 we observed effects on the inter-hemispheric tracts. Previous findings show that higher total white matter volume was associated with more social problems in individuals with NF1.^64^ Similarly, NF1 mice models with a biallelic deletion of *Nf1* in myelinating neuroglial progenitor cells show social impairments (less vocalizations).^65^ Increased RD and MD in the arcuate and the uncinate have also been reported in idiopathic autism spectrum disorders (ASD),^66^ but findings are mixed.^67^ These findings provide a more fine-grained picture of the effects of individual upstream Ras effectors on white-matter pathology and can assist in directing biomarker research for outcome measures in intervention studies and/or clinical trials.

Individuals with NS and NF1 often present mild-to-severe impairments in social cognition, communication, and interaction.^20,54,55^ In our independent cohort we observed elevated SRS scores in NS (p=.002) and NF1 (p<.001) compared to controls. ASD prevalence is also higher in these groups (25-30% compared to 1.7-2.3% in the general population).^54–56^ Abnormalities in white-matter microstructural properties of association tracts could account for the social deficits associated with these two groups. In this context, our findings elucidate condition-specific on FA special and magnitude effects and potential transdiagnostic therapeutic targets across Ras-MAPK pathway pathologies. That is, highly similar psychiatric outcomes (i.e., social deficits) may be partly driven by different underlying biological mechanisms. More broadly, with the increasing number of studies implicating Rasopathy-linked genes (i.e., *SYNGAP, SPRED1, NF1*) in the polygenic etiology of idiopathic autism spectrum disorders,^57–61^ studying the white-matter microstructural properties of association tracts in Rasopathies also has translational implications for idiopathic disorders.

This study is not without limitations. First, diffusion metrics are indirect indices of white-matter integrity as they measure water diffusion rather than biological substrate^68^. Therefore, discussions of underlying biological mechanisms are speculative and based on the histological findings from animal models of both diseases. Second, although anatomically constrained tractography tools like TRACULA are robust methods, crossing fibers remain challenging when computing diffusion metrics.^43^ Third, a modest sample size. We address this limitation by including effect sizes, which are independent of sample size and superior indicators of clinical significance^47^; a narrow age range of children, matching groups on sexual development (Table 1) with limited genetic diversity (restricting mutations in the NS group to *PTPN11* and *SOS1*) and capitalizing on the large effects genetic syndromes exert on brain morphometry.^69^ Finally, diffusion imaging is highly sensitive to motion and, despite incorporating advanced motion correction methods,^28–32,42^ data across all groups was discarded, likely excluding children with comorbid disorders prone to high motion. The results of this study should be interpreted considering these limitations.

In summary, our findings provide evidence of (1) pervasive effects of NS and NF1 on white matter integrity and (2) convergent effects of both syndromes on white matter integrity as evidenced by voxel-wise and tract-based analyses. These findings suggest the Ras-MAPK pathway is an important regulator of white-matter development in children; we also found evidence of (3) divergent spatial patterns and differences in magnitude effects between NS and NF1 on white matter integrity and of (4) largely affected and overlapping social phenotypes in and across clinical groups. With the increasing availability of pharmacological treatments targeting Ras-MAPK pathway hyperactivation for pediatric populations, this work is timely as it demonstrates the pervasive effects of Ras-MAPK hyperactivation on white-matter microstructure in children and illustrates potential condition-specific therapeutic targets.

## Supplementary material

Supplementary material is available online.

## Data availability statement

The deidentified data supporting the results of this study are available upon reasonable request to the corresponding author in compliance with applicable privacy laws, data protection and requirements for consent and anonymization.

## Acknowledgements

We thank all the children and their families who kindly volunteered to participate, the Noonan Syndrome Foundation, and the Children’s Tumor Foundation which made this work possible. We would also like to thank Dr. Jennifer Bruno (Stanford University), as well as current and past members of the BRIDGE Lab at the Psychiatry and Behavioral Sciences department at Stanford University for their support with data collection and intellectual discussions. Some of the computing for this project was performed on the Sherlock cluster. We would like to thank Stanford University and the Stanford Research Computing Center for providing computational resources and support that contributed to these research results.

## References

1. Boguski MS, McCormick F. Proteins regulating Ras and its relatives. Nature. 1993;366(6456):643–654. doi:10.1038/366643a0

2. Lavoie H, Gagnon J, Therrien M. ERK signalling: a master regulator of cell behaviour, life and fate. Nat Rev Mol Cell Biol. 2020;21(10):607–632. doi:10.1038/s41580-020-0255-7

3. Tidyman WE, Rauen KA. Pathogenetics of the RASopathies. Hum Mol Genet. 2016;25(R2):R123–R132. doi:10.1093/hmg/ddw191

4. Roberts AE, Allanson JE, Tartaglia M, Gelb BD. Noonan syndrome. Lancet. 2013;381(9863):333–342. doi:10.1016/S0140-6736(12)61023-X

5. Uusitalo E, Leppävirta J, Koffert A, et al. Incidence and mortality of neurofibromatosis: a total population study in Finland. J Invest Dermatol. 2015;135(3):904–906. doi:10.1038/jid.2014.465

6. Tartaglia M, Aoki Y, Gelb BD. The molecular genetics of RASopathies: An update on novel disease genes and new disorders. Am J Med Genet C Semin Med Genet. 2022;190(4):425–439. doi:10.1002/ajmg.c.32012

7. Wallace MR, Marchuk DA, Andersen LB, et al. Type 1 neurofibromatosis gene: identification of a large transcript disrupted in three NF1 patients. Science. 1990;249(4965):181-186. doi:10.1126/science.2134734

8. Ishii A, Furusho M, Dupree JL, Bansal R. Role of ERK1/2 MAPK signaling in the maintenance of myelin and axonal integrity in the adult CNS. J Neurosci. 2014;34(48):16031–16045. doi:10.1523/JNEUROSCI.3360-14.2014

9. Wahl SE, McLane LE, Bercury KK, Macklin WB, Wood TL. Mammalian target of rapamycin promotes oligodendrocyte differentiation, initiation and extent of CNS myelination. J Neurosci. 2014;34(13):4453–4465. doi:10.1523/JNEUROSCI.4311-13.2014

10. Ehrman LA, Nardini D, Ehrman S, et al. The protein tyrosine phosphatase Shp2 is required for the generation of oligodendrocyte progenitor cells and myelination in the mouse telencephalon. J Neurosci. 2014;34(10):3767–3778. doi:10.1523/JNEUROSCI.3515-13.2014

11. Asleh J, Shofty B, Cohen N, et al. Brain-wide structural and functional disruption in mice with oligodendrocyte-specific Nf1 deletion is rescued by inhibition of nitric oxide synthase. Proc Natl Acad Sci U S A. 2020;117(36):22506–22513. doi:10.1073/pnas.2008391117

12. Bennett MR, Rizvi TA, Karyala S, McKinnon RD, Ratner N. Aberrant growth and differentiation of oligodendrocyte progenitors in neurofibromatosis type 1 mutants. J Neurosci. 2003;23(18):7207–7217. doi:10.1523/JNEUROSCI.23-18-07207.2003

13. Fattah M, Raman MM, Reiss AL, Green T. PTPN11 Mutations in the Ras-MAPK Signaling Pathway Affect Human White Matter Microstructure. Cerebral Cortex. 2021;31(3):1489–1499. doi:10.1093/cercor/bhaa299

14. Karlsgodt KH, Rosser T, Lutkenhoff ES, Cannon TD, Silva A, Bearden CE. Alterations in white matter microstructure in neurofibromatosis-1. PLoS One. 2012;7(10):e47854. doi:10.1371/journal.pone.0047854

15. Wignall EL, Griffiths PD, Papadakis NG, et al. Corpus callosum morphology and microstructure assessed using structural MR imaging and diffusion tensor imaging: initial findings in adults with neurofibromatosis type 1. AJNR Am J Neuroradiol. 2010;31(5):856–861. http://www.ajnr.org/content/31/5/856.short

16. Aydin S, Kurtcan S, Alkan A, et al. Relationship between the corpus callosum and neurocognitive disabilities in children with NF-1: diffusion tensor imaging features. Clin Imaging. 2016;40(6):1092–1095. doi:10.1016/j.clinimag.2016.06.013

17. Filippi CG, Watts R, Duy LAN, Cauley KA. Diffusion-tensor imaging derived metrics of the corpus callosum in children with neurofibromatosis type I. AJR Am J Roentgenol. 2013;200(1):44–49. doi:10.2214/AJR.12.9590

18. Herbet G, Lafargue G, Bonnetblanc F, Moritz-Gasser S, de Champfleur NM, Duffau H. Inferring a dual-stream model of mentalizing from associative white matter fibres disconnection. Brain. 2014;137(3):944–959. doi:10.1093/brain/awt370

19. Parkinson C, Wheatley T. Relating anatomical and social connectivity: white matter microstructure predicts emotional empathy. Cereb Cortex. 2014;24(3):614–625. doi:10.1093/cercor/bhs347

20. Bruno J, Naylor P, Green T. Multidimensional Neuropsychiatric Phenotypes in Children With Noonan Syndrome. In: NEUROPSYCHOPHARMACOLOGY. Vol 47. SPRINGERNATURE CAMPUS, 4 CRINAN ST, LONDON, N1 9XW, ENGLAND; 2022:147–148.

21. Walsh KS, del Castillo A, Kennedy T, Karim AI, Semerjian C. A Review of Psychological, Social, and Behavioral Functions in the RASopathies. Journal of Pediatric Neuropsychology. 2020;6(3):131–142. doi:10.1007/s40817-020-00088-1

22. Marshall WA, Tanner JM. Variations in pattern of pubertal changes in girls. Arch Dis Child. 1969;44(235):291–303. doi:10.1136/adc.44.235.291

23. Marshall WA, Tanner JM. Variations in the pattern of pubertal changes in boys. Arch Dis Child. 1970;45(239):13–23. doi:10.1136/adc.45.239.13

24. Wechsler D. Wechsler Abbreviated Scale of Intelligence - Second Edition. Pearson; 2011. doi:10.1037/t15171-000

25. Constantino JN. Social Responsiveness Scale, Second Edition. Pearson; 2012.

26. Rai B, Naylor PE, Siqueiros-Sanchez M, et al. Novel effects of Ras-MAPK pathogenic variants on the developing human brain and their link to gene expression and inhibition abilities. Transl Psychiatry. 2023;13(1):245. doi:10.1038/s41398-023-02504-4

27. Siqueiros-Sanchez M, Rai B, Chowdhury S, Reiss AL, Green T. Syndrome specific neuroanatomical phenotypes in girls with Turner and Noonan Syndromes. Biol Psychiatry Cogn Neurosci Neuroimaging. Published online September 6, 2022. doi:10.1016/j.bpsc.2022.08.012

28. Andersson JLR, Skare S, Ashburner J. How to correct susceptibility distortions in spin-echo echo-planar images: application to diffusion tensor imaging. Neuroimage. 2003;20(2):870–888. doi:10.1016/S1053-8119(03)00336-7

29. Andersson JLR, Graham MS, Drobnjak I, Zhang H, Campbell J. Susceptibility-induced distortion that varies due to motion: Correction in diffusion MR without acquiring additional data. Neuroimage. 2018;171:277–295. doi:10.1016/j.neuroimage.2017.12.040

30. Andersson JLR, Graham MS, Drobnjak I, Zhang H, Filippini N, Bastiani M. Towards a comprehensive framework for movement and distortion correction of diffusion MR images: Within volume movement. Neuroimage. 2017;152:450–466. doi:10.1016/j.neuroimage.2017.02.085

31. Andersson JLR, Graham MS, Zsoldos E, Sotiropoulos SN. Incorporating outlier detection and replacement into a non-parametric framework for movement and distortion correction of diffusion MR images. Neuroimage. 2016;141:556–572. doi:10.1016/j.neuroimage.2016.06.058

32. Andersson JLR, Sotiropoulos SN. An integrated approach to correction for off-resonance effects and subject movement in diffusion MR imaging. Neuroimage. 2016;125:1063–1078. doi:10.1016/j.neuroimage.2015.10.019

33. Dale AM, Fischl B, Sereno MI. Cortical surface-based analysis. I. Segmentation and surface reconstruction. Neuroimage. 1999;9(2):179–194. doi:10.1006/nimg.1998.0395

34. Dale AM, Sereno MI. Improved localization of cortical activity by combining EEG and MEG with MRI cortical surface reconstruction: a linear approach. J Cogn Neurosci. 1993;5:162–176.

35. Fischl B, Dale AM. Measuring the thickness of the human cerebral cortex from magnetic resonance images. Proc Natl Acad Sci U S A. 2000;97(20):11050–11055. doi:10.1073/pnas.200033797

36. Fischl B, Salat DH, Busa E, et al. Whole brain segmentation: automated labeling of neuroanatomical structures in the human brain. Neuron. 2002;33(3):341–355. https://www.ncbi.nlm.nih.gov/pubmed/11832223

37. Fischl B, van der Kouwe A, Destrieux C, et al. Automatically parcellating the human cerebral cortex. Cereb Cortex. 2004;14(1):11–22. doi:10.1093/cercor/bhg087

38. Mukherjee P, Berman JI, Chung SW, Hess CP, Henry RG. Diffusion tensor MR imaging and fiber tractography: theoretic underpinnings. AJNR Am J Neuroradiol. 2008;29(4):632–641. doi:10.3174/ajnr.A1051

39. Larvie M, Fischl B. Volumetric and fiber-tracing MRI methods for gray and white matter. Handb Clin Neurol. 2016;135:39–60. doi:10.1016/B978-0-444-53485-9.00003-9

40. Song SK, Yoshino J, Le TQ, et al. Demyelination increases radial diffusivity in corpus callosum of mouse brain. Neuroimage. 2005;26(1):132–140. doi:10.1016/j.neuroimage.2005.01.028

41. Smith SM, Jenkinson M, Johansen-Berg H, et al. Tract-based spatial statistics: Voxelwise analysis of multi-subject diffusion data. Neuroimage. 2006;31(4):1487–1505. doi:10.1016/j.neuroimage.2006.02.024

42. Smith SM, Jenkinson M, Woolrich MW, et al. Advances in functional and structural MR image analysis and implementation as FSL. Neuroimage. 2004;23 Suppl 1:S208–19. doi:10.1016/j.neuroimage.2004.07.051

43. Maffei C, Lee C, Planich M, et al. Using diffusion MRI data acquired with ultra-high gradient strength to improve tractography in routine-quality data. Neuroimage. 2021;245:118706. doi:10.1016/j.neuroimage.2021.118706

44. Maffei C, Gilmore N, Snider SB, et al. Automated detection of axonal damage along white matter tracts in acute severe traumatic brain injury. NeuroImage Clin. 2023;37(103294):103294. doi:10.1016/j.nicl.2022.103294

45. Yendiki A, Koldewyn K, Kakunoori S, Kanwisher N, Fischl B. Spurious group differences due to head motion in a diffusion MRI study. Neuroimage. 2014;88:79–90. doi:10.1016/j.neuroimage.2013.11.027

46. Winkler AM, Ridgway GR, Webster MA, Smith SM, Nichols TE. Permutation inference for the general linear model. Neuroimage. 2014;92:381–397. doi:10.1016/j.neuroimage.2014.01.060

47. Kerby DS. The Simple Difference Formula: An Approach to Teaching Nonparametric Correlation. Comprehensive Psychology. 2014;3:11.IT.3.1. doi:10.2466/11.IT.3.1

48. de Blank P, Nishiyama A, López-Juárez A. A new era for myelin research in Neurofibromatosis type 1. Glia. Published online June 29, 2023. doi:10.1002/glia.24432

49. Wang Y, Kim E, Wang X, et al. ERK inhibition rescues defects in fate specification of Nf1-deficient neural progenitors and brain abnormalities. Cell. 2012;150(4):816–830. doi:10.1016/j.cell.2012.06.034

50. Mayes DA, Rizvi TA, Titus-Mitchell H, et al. Nf1 loss and Ras hyperactivation in oligodendrocytes induce NOS-driven defects in myelin and vasculature. Cell Rep. 2013;4(6):1197–1212. doi:10.1016/j.celrep.2013.08.011

51. Klawiter EC, Schmidt RE, Trinkaus K, et al. Radial diffusivity predicts demyelination in ex vivo multiple sclerosis spinal cords. Neuroimage. 2011;55(4):1454–1460. doi:10.1016/j.neuroimage.2011.01.007

52. López-Juárez A, Titus HE, Silbak SH, et al. Oligodendrocyte Nf1 Controls Aberrant Notch Activation and Regulates Myelin Structure and Behavior. Cell Rep. 2017;19(3):545–557. doi:10.1016/j.celrep.2017.03.073

53. Koini M, Rombouts SARB, Veer IM, Van Buchem MA, Huijbregts SCJ. White matter microstructure of patients with neurofibromatosis type 1 and its relation to inhibitory control. Brain Imaging Behav. 2017;11(6):1731–1740. doi:10.1007/s11682-016-9641-3

54. Garg S, Brooks A, Burns A, et al. Autism spectrum disorder and other neurobehavioural comorbidities in rare disorders of the Ras/MAPK pathway. Dev Med Child Neurol. 2017;59(5):544–549. doi:10.1111/dmcn.13394

55. Garg S, Lehtonen A, Huson SM, et al. Autism and other psychiatric comorbidity in neurofibromatosis type 1: evidence from a population-based study. Dev Med Child Neurol. 2013;55(2):139–145. doi:10.1111/dmcn.12043

56. Garg S, Green J, Leadbitter K, et al. Neurofibromatosis type 1 and autism spectrum disorder. Pediatrics. 2013;132(6):e1642–8. doi:10.1542/peds.2013-1868

57. Borrie SC, Brems H, Legius E, Bagni C. Cognitive Dysfunctions in Intellectual Disabilities: The Contributions of the Ras-MAPK and PI3K-AKT-mTOR Pathways. Annu Rev Genomics Hum Genet. 2017;18:115–142. doi:10.1146/annurev-genom-091416-035332

58. Borrie SC, Plasschaert E, Callaerts-Vegh Z, et al. MEK inhibition ameliorates social behavior phenotypes in a Spred1 knockout mouse model for RASopathy disorders. Mol Autism. 2021;12(1):53. doi:10.1186/s13229-021-00458-2

59. Pinto D, Delaby E, Merico D, et al. Convergence of genes and cellular pathways dysregulated in autism spectrum disorders. Am J Hum Genet. 2014;94(5):677–694. doi:10.1016/j.ajhg.2014.03.018

60. Pinto D, Pagnamenta AT, Klei L, et al. Functional impact of global rare copy number variation in autism spectrum disorders. Nature. 2010;466(7304):368–372. doi:10.1038/nature09146

61. Fu JM, Satterstrom FK, Peng M, et al. Rare coding variation provides insight into the genetic architecture and phenotypic context of autism. Nat Genet. 2022;54(9):1320–1331. doi:10.1038/s41588-022-01104-0

62. Catani M, Mesulam M. The arcuate fasciculus and the disconnection theme in language and aphasia: history and current state. Cortex. 2008;44(8):953–961. doi:10.1016/j.cortex.2008.04.002

63. Dick AS, Tremblay P. Beyond the arcuate fasciculus: consensus and controversy in the connectional anatomy of language. Brain. 2012;135(Pt 12):3529–3550. doi:10.1093/brain/aws222

64. Huijbregts SC, Loitfelder M, Rombouts SA, et al. Cerebral volumetric abnormalities in Neurofibromatosis type 1: associations with parent ratings of social and attention problems, executive dysfunction, and autistic mannerisms. J Neurodev Disord. 2015;7:32. doi:10.1186/s11689-015-9128-3

65. Maloney SE, Chandler KC, Anastasaki C, Rieger MA, Gutmann DH, Dougherty JD. Characterization of early communicative behavior in mouse models of neurofibromatosis type 1. Autism Res. 2018;11(1):44–58. doi:10.1002/aur.1853

66. Catani M, Dell’Acqua F, Budisavljevic S, et al. Frontal networks in adults with autism spectrum disorder. Brain. 2016;139(Pt 2):616–630. doi:10.1093/brain/awv351

67. Ameis SH, Catani M. Altered white matter connectivity as a neural substrate for social impairment in Autism Spectrum Disorder. Cortex. 2015;62:158–181. doi:10.1016/j.cortex.2014.10.014

68. Tournier JD, Mori S, Leemans A. Diffusion tensor imaging and beyond. Magn Reson Med. 2011;65(6):1532–1556. doi:10.1002/mrm.22924

69. Cheon EJ, Bearden CE, Sun D, et al. Cross disorder comparisons of brain structure in schizophrenia, bipolar disorder, major depressive disorder, and 22q11.2 deletion syndrome: A review of ENIGMA findings. Psychiatry Clin Neurosci. 2022;76(5):140–161. doi:10.1111/pcn.13337

