## Supplementary material for "Impact of Pathogenic Variants of the Ras-MAPK Pathway on Major White Matter Tracts in the Human Brain"

**Methods**

**Image acquisition, image preprocessing and quality control**

All participants completed behavioral training in a mock MRI scanner to familiarize themselves with the MRI environment and minimize motion-related artifacts during the following real MRI scan. Participants received feedback and guidance on their motion throughout this training scan. Participants were instructed to relax and remain still in the scanner during the training in the mock scanner and the real MRI scan.

*Diffusion weighted imaging.* Diffusion-weighted images (DWI) were collected using a single-shot diffusion-weighted spin-echo echo planar imaging sequence with the following parameters: field of view (FOV)=240 x 240 mm^2^; voxel size = 1.7 × 1.7 × 1.7 mm^3^; slice number= 75; multiband (MB) factor=3; no in-plane under-sampling (R=1); partial Fourier (F) factor=0.61 with Homodyne reconstruction^1^; repetition time (TR)=8300 ms; echo time (TE)=70.5 ms;  acquisition time= 15:05 min. Multi-shell diffusion encoding was played in a total of 96 directions with: *b*=500 s/mm^2^ (6-dirs); *b*=1000 s/mm^2^ (15-dirs); *b*=2000 s/mm^2^ (15-dirs); *b*=3000 s/mm^2^ (60-dirs). The sequence included one b=0 scan was interspersed between each 10 diffusion volumes for subject bulk motion correction (6 “blip up”). Three extra b=0 datasets were acquired with reversed phase encoding (PE) polarity (3 “blip down”) for distortion correction. Preprocessing of DWI included correcting for susceptibility-induced distortions using FSL’s topup ^2^. Corrections for eddy current distortions and motion artifacts were done with FSL’s eddy^3–6^ using the slice-to-volume motion model, susceptibility-by-movement correction, and the replace outliers options. Independent raters (MS, MR) examined the corrected DWI output. Nine subjects (5-NF1, 2-NS, 2-TD) were removed for not meeting minimum quality for pre-processing (n=5) or after preprocessing (n=4).

*T1 weighted imaging.* Whole-brain high-resolution T1-weighted (T1w) magnetization-prepared rapid gradient-echo (MPRAGE) images were collected during the same session, with the following main parameters: TR=7ms; TE = 2.83 ms, inversion time (TI) = 900 ms, flip angle = 8°, number of excitations (NEX) = 1; field of view (FOV) = 240 × 240 × 180 mm^3^; voxel size = 1.0 × 1.0 × 1.2 mm^3^; acquisition time = 4:22 min. T1 weighted images were preprocessed with the standard Freesurfer 7.2.0 recon-all pipeline (https://surfer.nmr.mgh.harvard.edu). After Freesurfer reconstruction and segmentation, independent raters (MR, CM) inspected. Edits were performed by trained personnel (CM) and the freesurfer pipeline was re-run until the surfaces for each T1 met quality standards for 2 independent raters (MR, CM).

**Head motion**

To assess for head motion differences, groups were compared on their Euler numbers from T1 structural imaging, mean diffusion signal-to-noise ratios and Eddy average translation. There were no differences in head motion between the groups in any of the measures: Euler number (F(2)=1.014,	p=0.369); Mean diffusion signal-to-noise ratio (F(2)=0.804, p=0.452); Eddy average translation (F(2)=2.067, p=0.136).

**Tract-Based Spatial Statistics (TBSS)**

First, FA, AD, RD, and MD images were created by fitting a diffusion tensor model to the raw diffusion data using FMRIB's Diffusion Toolbox^7^. Individual FA images were then non-linearly registered and transformed to the FSL standard space 1x1x1mm FMRIB58 FA template (<https://fsl.fmrib.ox.ac.uk/fsl/fslwiki/FMRIB58_FA>)^8,9^. Next, the mean FA image was created and thinned to create a mean FA skeleton using a threshold of 0.2. This skeleton represents the center of all tracts common to all subjects. Each subject’s aligned FA data was then projected onto the FA skeleton. AD, RD, and MD images were processed in the same manner (using the transformations obtained from the FA images and mapped onto the FA skeleton). The resulting skeletonized FA, AD, RD, and MD aligned data was used to calculate voxel-wise between-group statistics.

**TRActs Constrained by UnderLying Anatomy (TRACULA)**

In TRACULA, the diffusion scalars are quantified from anatomically corresponding regions (the major white matter tracts) on a subject-per-subject basis. TRACULA combines distortion-corrected diffusion data (fitting a “ball-and-stick” model at every voxel using FSL’s BEDPOSTX tool) data with T1w image-derived anatomical neighborhood priors (generated using the FreeSurfer recon-all pipeline) to reconstruct white matter tracts for each subject. To establish the anatomical priors for each subject, the TRACULA algorithm incorporates a priori anatomical knowledge derived from a set of individuals whose white-matter tracts were manually annotated and used as a training set ^10,11^ and combines it with the subject’s T1w data.

**Quality control and excluded tracts per group.** Only successfully reconstructed tracts were included (Table S1). According to Maffei et al ^12^, a failed reconstruction happens when probabilistic tractography is not able to find an appropriate reconstruction solution for the whole or a part of the tract. All 3D-reconstructions and volumes were visually inspected (MS) to identify failed and partial reconstructions of individual tracts as described in Maffei et al. Tracts that failed or were partially reconstructed were rerun using TRACULA’s built-in *reinit* function which uses a different set of initial points to reconstruct the tract. Tracts that failed to reconstruct or did so partially after *reinit* were excluded (Table S1). Tracts excluded based on quality were the anterior commissure, the acoustic radiation, the ventral cingulum bundle, and the fornix.

**PASTA analyses**

PASTA analyses are conducted on both sides of a tract located on both hemispheres (left and right). Participants where either the left or right tract has been excluded at the whole-tract level analysis due to quality are excluded from the PASTA analysis of said tract.

### Table S1. Number of tracts analyzed and excluded in TRACULA and analyzed in PASTA per group

|  | **TRACULA** | | | **PASTA** | | |
| --- | --- | --- | --- | --- | --- | --- |
|  | NS  (*n*= 22) | NF1  (*n=* 19) | TD  (*n=* 26) | NS  (*n*= 22) | NF1  (*n=* 19) | TD  (*n=* 26) |
| **Analyzed Tracts** |  |  |  |  |  |  |
| CC body |  |  |  |  |  |  |
| central | 2 | 1 | 0 | - | - | - |
| parietal | 0 | 0 | 0 | - | - | - |
| prefrontal | 0 | 0 | 0 | - | - | - |
| premotor | 0 | 0 | 0 | - | - | - |
| temporal | 0 | 1 | 0 | - | - | - |
| CC genu | 0 | 0 | 1 | - | - | - |
| CC rostrum | 5 | 1 | 3 | - | - | - |
| CC splenium | 0 | 0 | 1 | - | - | - |
| Left ATR | 0 | 0 | 0 | - | - | - |
| Right ATR | 1 | 0 | 1 | - | - | - |
| Left AF | 0 | 1 | 1 | 22 | 17 | 25 |
| Right AF | 0 | 0 | 0 |  |  |  |
| Left FAT | 0 | 0 | 0 | 22 | 18 | 26 |
| Right FAT | 0 | 0 | 0 |  |  |  |
| Left ILF | 0 | 0 | 1 | 21 | 18 | 25 |
| Right ILF | 1 | 0 | 0 |  |  |  |
| Left MLF | 1 | 2 | 0 | 21 | 16 | 26 |
| Right MLF | 1 | 1 | 0 |  |  |  |
| Left SLF I | 0 | 0 | 0 | 22 | 18 | 26 |
| Right SLF I | 0 | 0 | 0 |  |  |  |
| Left SLF II | 0 | 0 | 0 | 22 | 18 | 26 |
| Right SLF II | 0 | 0 | 0 |  |  |  |
| Left SLF III | 0 | 0 | 0 | 22 | 18 | 26 |
| Right SLF III | 0 | 0 | 0 |  |  |  |
| Left UF | 3 | 0 | 3 | 19 | 18 | 23 |
| Right UF | 2 | 0 | 2 |  |  |  |
| **Excluded Tracts** |  |  |  |  |  |  |
| Anterior commissure | 16 | 11 | 15 | - | - | - |
| Left acoustic radiation | 6 | 2 | 2 | - | - | - |
| Right acoustic radiation | 4 | 0 | 2 | - | - | - |
| Left cingulum bundle dorsal | 1 | 1 | 0 | - | - | - |
| Right cingulum bundle dorsal | 1 | 0 | 0 | - | - | - |
| Left cingulum bundle ventral | 7 | 4 | 6 | - | - | - |
| Left corticospinal tract | 0 | 1 | 0 | - | - | - |
| Right corticospinal tract | 0 | 1 | 0 | - | - | - |
| Mid cerebellar peduncle | 0 | 0 | 0 | - | - | - |
| Left external capsule | 4 | 0 | 2 | - | - | - |
| Right external capsule | 6 | 1 | 4 | - | - | - |
| Left fornix | 7 | 8 | 4 | - | - | - |
| Right fornix | 9 | 5 | 6 | - | - | - |
| Left optic radiation | 1 | 0 | 0 | - | - | - |
| Right optic radiation | 1 | 0 | 2 | - | - | - |

AF: Arcuate fasciculus; ATR: Anterior thalamic radiation; CC: Corpus callosum; FAT: Frontal aslant tract; ILF: Inferior longitudinal fasciculus; MLF: Middle longitudinal fasciculus; SLF: Superior longitudinal fasciculus; UF: Uncinate fasciculus.
Table S2. Descriptive statistics of individual tracts FA per group

|  | **Tract** | **NS** | | **NF1** | | **TD** | |  | **NS-TD** | |  | **NF1-TD** | | **NS-NF1** | | |
| --- | --- | --- | --- | --- | --- | --- | --- | --- | --- | --- | --- | --- | --- | --- | --- | --- |
|  |  | Mean (SD) | | Mean (SD) | | Mean (SD) | | *d* | 95% CI | | *d* | 95% CI | | *d* | 95% CI | |
| FA | CC body |  |  |  |  |  |  |  |  |  |  |  |  |  |  | |
|  | central | .561 (.033) | | .569 (.036) | | .588 (.022) | | -0.97 | -0.35, -1.6 | | -0.67 | -0.04, -1.3 | | -0.22 | -0.87, 0.43 | |
|  | parietal | .571 (.023) | | .561 (.031) | | .591 (.027) | | -0.81 | -0.21, -1.4 | | -1.07 | -0.42, -1.7 | | 0.38 | -0.25, 1.01 | |
|  | prefrontal | .560 (.032) | | .545 (.028) | | .572 (.027) | | -0.41 | 0.17, -0.98 | | -0.98 | -0.34, -1.61 | | 0.49 | -0.14, 1.12 | |
|  | premotor | .599 (.022) | | .586 (.024) | | .612 (.026) | | -0.52 | 0.06, -1.09 | | -1.01 | -0.37, -1.64 | | 0.57 | -0.07, 1.21 | |
|  | temporal | .521 (.022) | | .533 (.027) | | .551 (.019) | | -1.45 | -0.80, -2.08 | | -0.80 | -0.16, -1.43 | | -0.47 | -1.11, 0.17 | |
|  | CC genu | .565 (.034) | | .552 (.027) | | .582 (.022) | | -0.60 | -0.01, -1.18 | | -1.24 | -0.57, -1.89 | | 0.43 | -0.21, 1.06 | |
|  | CC rostrum | .543 (.059) | | .554 (.023) | | .566 (.045) | | -0.44 | 0.20, -1.07 | | -0.32 | 0.31, -0.95 | | -0.23 | -0.91, 0.44 | |
|  | CC splenium | .620 (.034) | | .618 (.043) | | .655 (.038) | | -0.99 | -0.37, -1.59 | | -0.94 | -0.29, -1.57 | | 0.05 | -0.57, 0.68 | |
|  | L ATR | .454(.020) | | .459 (.028) | | .484 (.021) | | -1.44 | -0.80, -2.08 | | -1.03 | -0.39, -1.67 | | -0.22 | -0.84, 0.41 | |
|  | R ATR | .441(.023) | | .439 (.017) | | .469 (.022) | | -1.24 | -0.60, -1.87 | | -1.49 | -0.8, -2.17 | | 0.11 | -0.52, 0.74 | |
|  | L AF | .443 (.021) | | .466 (.031) | | .473 (.023) | | -1.37 | -0.73, -2 | | -0.24 | 0.38, -0.85 | | -0.92 | -1.58, -0.25 | |
|  | R AF | .426(.021) | | .453 (.028) | | .456 (.024) | | -1.29 | -0.66, -1.91 | | -0.12 | 0.48, -0.72 | | -1.08 | -1.74, -0.41 | |
|  | L FAT | .487 (.030) | | .495 (.033) | | .502 (.037) | | -0.47 | 0.11, -1.04 | | -0.21 | 0.39, -0.82 | | -0.26 | -0.89, 0.37 | |
|  | R FAT | .475(.024) | | .489 (.032) | | .501 (.029) | | -0.96 | -0.35, -1.55 | | -0.38 | 0.23, -0.99 | | -0.5 | -1.13, 0.14 | |
|  | L ILF | .493(.027) | | .504 (.030) | | .532 (.022) | | -1.59 | -0.92, -2.24 | | -1.09 | -0.43, -1.73 | | -0.38 | -1.01, 0.25 | |
|  | R ILF | .484(.022) | | .497 (.033) | | .527 (.028) | | -1.67 | -1, -2.33 | | -0.96 | -0.33, -1.61 | | -0.47 | -1.11, 0.17 | |
|  | L MLF | .470 (.026) | | .489 (.024) | | .512 (.021) | | -1.79 | -1.10, -2.47 | | -1.07 | -0.4, -1.73 | | -0.73 | -1.40, -0.06 | |
|  | R MLF | .453 (.019) | | .481 (.026) | | .494 (.024) | | -1.88 | -1.18, -2.57 | | -0.53 | 0.1, -1.14 | | -1.27 | -1.97, -0.56 | |
|  | L SLF I | .531 (.026) | | .532 (.032) | | .551 (.028) | | -0.76 | -0.17, -1.34 | | -0.63 | -0.01, -1.25 | | -0.06 | -0.68, 0.57 | |
|  | R SLF I | .530 (.025) | | .541 (.035) | | .558 (.026) | | -1.08 | -0.47, -1.68 | | -0.58 | 0.04, -1.19 | | -0.34 | -0.96, 0.29 | |
|  | L SLF II | .407 (.022) | | .428 (.030) | | .436 (.028) | | -1.15 | -0.53, -1.76 | | -0.28 | 0.33, -0.88 | | -0.81 | -1.46, -0.16 | |
|  | R SLF II | .409 (.023) | | .430 (.031) | | .434 (.028) | | -0.98 | -0.38, -1.58 | | -0.13 | 0.47, -0.73 | | -0.81 | -1.45, -0.15 | |
|  | L SLF III | .439 (.033) | | .462 (.029) | | .472 (.025) | | -1.14 | -0.52, -1.75 | | -0.38 | 0.23, -0.98 | | -0.74 | -1.38, -0.09 | |
|  | R SLF III | .418(.026) | | .441 (.031) | | .451 (.027) | | -1.22 | -0.59, -1.83 | | -0.35 | 0.26, -0.95 | | -0.80 | -1.45, -0.15 | |
|  | L UF | .443(.026) | | .455 (.031) | | .472 (.031) | | -1.03 | -0.38, -1.67 | | -0.55 | 0.08, -1.18 | | -0.44 | -1.08, 0.22 | |
|  | R UF | .445 (.032) | | .467 (.027) | | .477 (.039) | | -0.88 | -0.25, -1.5 | | -0.30 | 0.31, -0.92 | | -0.71 | -1.37, -0.05 | |
| AD | CC body |  | |  | |  | |  |  | |  |  | |  |  | |
|  | central | .00092 (3.27E-05) | | .00093 (3.98E-05) | | .00091 (3.19E-05) | | 0.52 | -0.08, 1.11 | | 0.62 | -0.01, 1.24 | | -0.14 | 0.79, 0.51 | |
|  | parietal | .00094 (2.56E-05) | | .00095 (2.91E-05) | | .00092 (2.76E-05) | | 0.78 | 0.19, 1.37 | | 0.86 | 0.23, 1.48 | | -0.12 | 0.75, 0.50 | |
|  | prefrontal | .00093 (2.96E-05) | | .00092 (3.44E-05) | | .00091 (3.26E-05) | | 0.64 | 0.06, 1.22 | | 0.38 | -0.23, 0.99 | | 0.23 | -0.4, 0.85 | |
|  | premotor | .00096 (2.53E-05) | | .00095 (3.24E-05) | | .00093 (2.69E-05) | | 1 | 0.39, 1.6 | | 0.69 | 0.06, 1.30 | | 0.21 | -0.42, 0.83 | |
|  | temporal | .00094 (2.75E-05) | | .00095 (2.78E-05) | | .00092 (2.68E-05) | | 0.89 | 0.29, 1.49 | | 1.24 | 0.56, 1.9 | | -0.34 | -0.98, 0.3 | |
|  | CC genu | .00096 (3.56E-05) | | .00096 (3.46E-05) | | .00095 (3.47E-05) | | 0.29 | -0.29, 0.87 | | 0.30 | -0.32, 0.91 | | -0.01 | -0.63, 0.62 | |
|  | CC rostrum | .00096 (3.43E-05) | | .00097 (3.97E-05) | | .00095 (3.91E-05) | | 0.32 | -0.31, 0.95 | | 0.51 | -0.13, 1.14 | | -0.21 | -0.88, 0.46 | |
|  | CC splenium | .001 (3.75E-05) | | .001 (3.82E-05) | | .00099 (3.28E-05) | | 0.01 | -0.56, 0.59 | | 0.15 | -0.46, 0.75 | | -0.12 | -0.75, 0.50 | |
|  | L ATR | .00084 (2.53E-05) | | .00087 (4.35E-05) | | .00083 (2.57E-05) | | 0.53 | -0.05, 1.10 | | 1.26 | 0.6, 1.92 | | -0.85 | -1.5, -0.19 | |
|  | R ATR | .00083 (2.35E-05) | | .00085 (3.78E-05) | | .00082 (2.42E-05) | | 0.56 | -0.04, 1.14 | | 1.04 | 0.39, 1.69 | | -0.60 | -1.24, 0.04 | |
|  | L AF | .00081 (3.07E-05) | | .00083 (3.46E-05) | | .00079 (2.54E-05) | | 0.77 | 0.17, 1.36 | | 1.50 | 0.8, 2.19 | | -0.7 | -1.35, -0.04 | |
|  | R AF | .00079 (3.61E-05) | | .00081 (2.96E-05) | | .00076 (2.66E-05) | | 0.73 | 0.14, 1.31 | | 1.69 | 0.98, 2.38 | | -0.73 | -1.37, -0.08 | |
|  | L FAT | .00083 (2.72E-05) | | .00084 (3.75E-05) | | .00080 (3.31E-05) | | 1.16 | 0.54, 1.77 | | 1.25 | 0.59, 1.90 | | -0.26 | -0.88, 0.37 | |
|  | R FAT | .00081 (2.75E-05) | | .00083 (3.49E-05) | | .00079 (3.24E-05) | | 0.88 | 0.28, 1.47 | | 1.23 | 0.56, 1.87 | | -0.47 | -1.1, 0.17 | |
|  | L ILF | .00095 (3.09E-05) | | .00096 (2.82E-05) | | .00092 (2.91E-05) | | 0.92 | 0.31, 1.51 | | 1.41 | 0.72, 2.08 | | -0.44 | -1.07, 0.19 | |
|  | R ILF | .00093 (3.34E-05) | | .00094 (2.30E-05) | | .00091 (3.24E-05) | | 0.84 | 0.23, 1.43 | | 1.06 | 0.41, 1.69 | | -0.11 | -0.74, 0.52 | |
|  | L MLF | .00093 (3.28E-05) | | .00094 (2.24E-05) | | .00091 (2.62E-05) | | 0.45 | -0.13, 1.04 | | 1.21 | 0.52, 1.88 | | -0.57 | -1.24, 0.09 | |
|  | R MLF | .0009 (2.84E-05) | | .00092 (2.67E-05) | | .00088 (2.48E-05) | | 0.85 | 0.25, 1.45 | | 1.56 | 0.85, 2.25 | | -0.62 | -1.27, 0.04 | |
|  | L SLF I | .00087 (3.66E-05) | | .00087 (3.52E-05) | | .00083 (3.00E-05) | | 1 | 0.39, 1.59 | | 1.13 | 0.47, 1.77 | | -0.09 | -0.71, 0.54 | |
|  | R SLF I | .00087 (2.69E-05) | | .00089 (3.91E-05) | | .00085 (2.38E-05) | | 0.85 | 0.26, 1.44 | | 1.12 | 0.47, 1.76 | | -0.4 | -1.03, 0.23 | |
|  | L SLF II | .00076 (3.11E-05) | | .00079 (3.04E-05) | | .00074 (2.63E-05) | | 0.67 | 0.08, 1.25 | | 1.49 | 0.8, 2.16 | | -0.73 | -1.37, -0.08 | |
|  | R SLF II | .00076 (3.53E-05) | | .00079 (2.50E-05) | | .00074 (2.52E-05) | | 0.72 | 0.13, 1.31 | | 1.86 | 1.13, 2.57 | | -0.79 | -1.44, -0.14 | |
|  | L SLF III | .00076 (2.99E-05) | | .00078 (3.19E-05) | | .00074 (2.11E-05) | | 1.01 | 0.40, 1.61 | | 1.61 | 0.91, 2.3 | | -0.52 | -1.15, 0.12 | |
|  | R SLF III | .00074 (3.08E-05) | | .00076 (2.98E-05) | | .00071 (2.32E-05) | | 0.92 | 0.32, 1.51 | | 1.75 | 1.03, 2.45 | | -0.68 | -1.32, -0.04 | |
|  | L UF | .00091 (2.60E-05) | | .00092 (2.41E-05) | | .00089 (3.10E-05) | | 0.52 | -0.1, 1.14 | | 0.84 | 0.19, 1.47 | | -0.34 | -0.99, 0.31 | |
|  | R UF | .00092 (2.43E-05) | | .00093 (3.21E-05) | | .00090 (3.12E-05) | | 0.57 | -0.04, 1.17 | | 0.9 | 0.25, 1.54 | | -0.44 | -1.08, 0.21 | |
| RD | CC body |  | |  | |  | |  |  | |  |  | |  | |  |
|  | central | .00033 (.00003) | | .00033 (.00003) | | .00030 (.00001) | | 1.49 | 0.82, 2.14 | | 1.28 | 0.6, 1.94 | | 0.09 | | -0.56, 0.73 |
|  | parietal | .00034 (.00002) | | .00035 (.00003) | | .00031 (.00002) | | 1.13 | 0.51, 1.73 | | 1.46 | 0.78, 2.13 | | -0.52 | | -1.16, 0.11 |
|  | prefrontal | .00034 (.00003) | | .00035 (.00003) | | .00032 (.00002) | | 0.93 | 0.33, 1.53 | | 1.36 | 0.69, 2.02 | | -0.43 | | -1.06, 0.2 |
|  | premotor | .00031 (.00002) | | .00033 (.00003) | | .00029 (.00002) | | 1.14 | 0.52, 1.74 | | 1.45 | 0.76, 2.11 | | -0.47 | | -1.09, 0.17 |
|  | temporal | .00038 (.00002) | | .00037 (.00003) | | .00034 (.00002) | | 1.89 | 1.2, 2.57 | | 1.45 | 0.76, 2.13 | | 0.17 | | -0.46, 0.81 |
|  | CC genu | .00034 (.00003) | | .00035 (.00003) | | .00032 (.00002) | | 0.96 | 0.35, 1.56 | | 1.42 | 0.73, 2.09 | | -0.41 | | -1.03, 0.23 |
|  | CC rostrum | .00037 (.00005) | | .00036 (.00003) | | .00034 (.00003) | | 0.64 | -0.01, 1.28 | | 0.66 | 0.01, 1.3 | | 0.15 | | -0.53, 0.82 |
|  | CC splenium | .00031 (.00002) | | .00032 (.00004) | | .00028 (.00003) | | 1.16 | 0.54, 1.78 | | 1.16 | 0.5, 1.81 | | -0.22 | | -0.84, 0.41 |
|  | L ATR | .00039 (.00002) | | .00040 (.00003) | | .00036 (.00002) | | 1.65 | 0.99, 2.31 | | 1.68 | 0.97, 2.37 | | -0.36 | | -0.99, 0.27 |
|  | R ATR | .00040 (.00002) | | .00041 (.00002) | | .00037 (.00001) | | 1.82 | 1.12, 2.5 | | 2.34 | 3.12, 1.54 | | -0.51 | | -1.15, 0.13 |
|  | L AF | .00039 (.00002) | | .00038 (.00003) | | .00035 (.00002) | | 1.76 | 1.08, 2.43 | | 1.14 | 0.47, 1.8 | | 0.32 | | -0.32, 0.96 |
|  | R AF | .00039 (.00002) | | .00038 (.00003) | | .00035 (.00002) | | 1.72 | 1.04, 2.38 | | 1.1 | 0.45, 1.74 | | 0.35 | | -0.28, 0.97 |
|  | L FAT | .00037 (.00002) | | .00037 (.00003) | | .00034 (.00002) | | 1.11 | 0.49, 1.71 | | 0.96 | 0.32, 1.59 | | 0.01 | | -0.61, 0.63 |
|  | R FAT | .00037 (.00002) | | .00036 (.00003) | | .00034 (.00002) | | 1.57 | 0.91, 2.22 | | 1.17 | 0.51, 1.81 | | 0.11 | | -0.51, 0.74 |
|  | L ILF | .00041 (.00003) | | .00041 (.00003) | | .00037 (.00002) | | 1.97 | 1.26, 2.67 | | 1.65 | 0.94, 2.34 | | 0.17 | | -0.46, 0.79 |
|  | R ILF | .00041 (.00002) | | .00040 (.00003) | | .00037 (.00002) | | 2.06 | 1.34, 2.77 | | 1.39 | 0.72, 2.06 | | 0.37 | | -0.27, 1 |
|  | L MLF | .00042 (.00002) | | .00041 (.00002) | | .00038 (.00002) | | 1.97 | 1.26, 2.66 | | 1.62 | 0.89, 2.32 | | 0.35 | | -0.31, 1.01 |
|  | R MLF | .00043 (.00002) | | .00041 (.00003) | | .00038 (.00002) | | 2.21 | 1.47, 2.94 | | 1.14 | 0.47, 1.79 | | 0.81 | | 0.14, 1.47 |
|  | L SLF I | .00035 (.00002) | | .00035 (.00003) | | .00032 (.00002) | | 1.46 | 0.82, 2.1 | | 1.28 | 0.62, 1.94 | | -0.09 | | -0.71, 0.53 |
|  | R SLF I | .00035 (.00002) | | .00035 (.00003) | | .00032 (.00002) | | 1.53 | 0.88, 2.18 | | 1.22 | 0.56, 1.87 | | 0.08 | | -0.55, 0.7 |
|  | L SLF II | .00040 (.00002) | | .00039 (.00003) | | .00036 (.00002) | | 1.55 | 0.9, 2.2 | | 1.15 | 0.49, 1.79 | | 0.2 | | -0.43, 0.82 |
|  | R SLF II | .00039 (.00002) | | .00039 (.00003) | | .00036 (.00002) | | 1.47 | 0.82, 2.1 | | 1.08 | 0.43, 1.72 | | 0.17 | -0.45, 0.79 | |
|  | L SLF III | .00038 (.00003) | | .00037 (.00003) | | .00034 (.00002) | | 1.56 | 0.91, 2.21 | | 1.15 | 0.49, 1.79 | | 0.38 | -0.25, 1.01 | |
|  | R SLF III | .00038 (.00002) | | .00037 (.00003) | | .00034 (.00002) | | 1.62 | 0.96, 2.27 | | 1.2 | 0.54, 1.85 | | 0.25 | -0.38, 0.87 | |
|  | L UF | .00043 (.00003) | | .00043 (.00003) | | .00040 (.00002) | | 1.37 | 0.68, 2.04 | | 0.95 | 0.3, 1.6 | | 0.21 | -0.44, 0.85 | |
|  | R UF | .00044 (.00003) | | .00042 (.00003) | | .00040 (.00003) | | 1.15 | 0.50, 1.79 | | 0.71 | 0.08, 1.34 | | 0.44 | -0.21, 1.08 | |
| MD | CC body |  | |  | |  | |  |  | |  |  | |  |  | |
|  | central | .00053 (.000020) | | .00053 (.000025) | | .0005 (.000014) | | 1.51 | 0.84, 2.17 | | 1.31 | 0.63, 1.97 | | -0.01 | -0.65, 0.64 | |
|  | parietal | .00054 (.000017) | | .00055 (.000025) | | .00052 (.000017) | | 1.34 | 0.7, 1.96 | | 1.55 | 0.86, 2.23 | | -0.47 | -1.1, 0.17 | |
|  | prefrontal | .00053 (.000021) | | .00054 (.00003) | | .00051 (.000014) | | 1.16 | 0.54, 1.77 | | 1.18 | 0.52, 1.83 | | -0.23 | -0.86, 0.4 | |
|  | premotor | .00053 (.000018) | | .00053 (.000028) | | .00051 (.000013) | | 1.47 | 0.82, 2.1 | | 1.38 | 0.71, 2.04 | | -0.23 | -0.86, 0.39 | |
|  | temporal | .00057 (.000018) | | .00057 (.000023) | | .00054 (.000015) | | 1.82 | 1.14, 2.49 | | 1.66 | 0.94, 2.36 | | -0.03 | -0.66, 0.61 | |
|  | CC genu | .00055 (.000022) | | .00056 (.00003) | | .00053 (.000017) | | 0.91 | 0.31, 1.51 | | 1.12 | 0.46, 1.76 | | -0.3 | -0.93, 0.32 | |
|  | CC rostrum | .00057 (.00003) | | .00056 (.000028) | | .00055 (.00002) | | 0.84 | 0.18, 1.49 | | 0.83 | 0.17, 1.48 | | 0.04 | -0.63, 0.71 | |
|  | CC splenium | .00054 (.000016) | | .00055 (.000026) | | .00052 (.000018) | | 1.14 | 0.51, 1.75 | | 1.16 | 0.5, 1.81 | | -0.27 | -0.9, 0.36 | |
|  | L ATR | .00054 (.000019) | | .00056 (.000031) | | .00052 (.000017) | | 1.4 | 0.76, 2.03 | | 1.72 | 1.01, 2.41 | | -0.63 | -1.26, 0.02 | |
|  | R ATR | .00054 (.000018) | | .00056 (.000027) | | .00052 (.00001) | | 1.67 | 0.98, 2.34 | | 2.01 | 1.25, 2.74 | | -0.61 | -1.25, 0.04 | |
|  | L AF | .00053 (.000022) | | .00053 (.000026) | | .0005 (.000015) | | 1.55 | 0.89, 2.2 | | 1.54 | 0.83, 2.23 | | -0.1 | -0.73, 0.54 | |
|  | R AF | .00052 (.000024) | | .00052 (.000027) | | .00049 (.000016) | | 1.48 | 0.83, 2.11 | | 1.53 | 0.84, 2.21 | | -0.09 | -0.72, 0.53 | |
|  | L FAT | .00052 (.000021) | | .00052 (.00003) | | .00049 (.000017) | | 1.52 | 0.87, 2.17 | | 1.37 | 0.69, 2.03 | | -0.1 | -0.72, 0.52 | |
|  | R FAT | .00052 (.00002) | | .00052 (.000028) | | .00049 (.000015) | | 1.63 | 0.97, 2.28 | | 1.48 | 0.8, 2.16 | | -0.12 | -0.75, 0.51 | |
|  | L ILF | .00059 (.000026) | | .00059 (.000028) | | .00055 (.000017) | | 1.85 | 1.15, 2.53 | | 1.85 | 1.12, 2.57 | | -0.04 | -0.66, 0.58 | |
|  | R ILF | .00059 (.000024) | | .00058 (.000026) | | .00055 (.00002) | | 1.87 | 1.17, 2.56 | | 1.56 | 0.86, 2.24 | | 0.23 | -0.4, 0.86 | |
|  | L MLF | .00059 (.000024) | | .00059 (.000021) | | .00056 (.000017) | | 1.6 | 0.93, 2.26 | | 1.73 | 1, 2.45 | | 0.01 | -0.64, 0.66 | |
|  | R MLF | .00059 (.000019) | | .00058 (.000024) | | .00055 (.000018) | | 1.99 | 1.28, 2.69 | | 1.50 | 0.8, 2.19 | | 0.3 | -0.35, 0.94 | |
|  | L SLF I | .00052 (.00002) | | .00053 (.000026) | | .00049 (.000014) | | 1.74 | 1.06, 2.4 | | 1.59 | 0.9, 2.28 | | -0.11 | -0.74, 0.51 | |
|  | R SLF I | .00053 (.000017) | | .00053 (.000024) | | .00050 (.000015) | | 1.68 | 1.01, 2.34 | | 1.56 | 0.86, 2.23 | | -0.16 | -0.78, 0.47 | |
|  | L SLF II | .00052 (.000022) | | .00052 (.000025) | | .00049 (.000017) | | 1.39 | 0.75, 2.02 | | 1.53 | 0.84, 2.21 | | -0.18 | -0.81, 0.44 | |
|  | R SLF II | .00052 (.000024) | | .00052 (.000025) | | .00049 (.000015) | | 1.37 | 0.73, 2 | | 1.65 | 0.95, 2.34 | | -0.23 | -0.85, 0.40 | |
|  | L SLF III | .0005 (.000023) | | .00050 (.000024) | | .00047 (.000015) | | 1.64 | 0.98, 2.3 | | 1.57 | 0.87, 2.25 | | 0.05 | -0.57, 0.67 | |
|  | R SLF III | .0005 (.000023) | | .00050 (.000026) | | .00047 (.000014) | | 1.59 | 0.93, 2.24 | | 1.68 | 0.97, 2.37 | | -0.11 | -0.73, 0.51 | |
|  | L UF | .00059 (.000022) | | .00059 (.000027) | | .00057 (.000016) | | 1.35 | 0.67, 2.02 | | 1.14 | 0.47, 1.8 | | 0.05 | -0.6, 0.69 | |
|  | R UF | .0006 (.000021) | | .00059 (.000027) | | .00057 (.000022) | | 1.28 | 0.62, 1.93 | | 0.98 | 0.33, 1.62 | | 0.17 | -0.47, 0.81 | |

###### Table S3. Between-group differences in tract-based mean fractional anisotropy, axial diffusivity, radial diffusivity and mean diffusivity of commissural, projection and association tracts.

| **Tract name** | | | **FA** | | | | **AD** | | | | **RD** | | | | **MD** | | | |
| --- | --- | --- | --- | --- | --- | --- | --- | --- | --- | --- | --- | --- | --- | --- | --- | --- | --- | --- |
|  | **ANCOVA**  ***p-*value*** | | | | ***t-*test p*-*value** | | **ANCOVA**  ***p-*value*** | | ***t-*test p*-*value** | | **ANCOVA**  ***p-*value*** | | ***t-*test p*-*value** | | **ANCOVA**  ***p-*value*** | | ***t-*test p*-*value** | |
|  |  | | | **NS-TD** | | **NF1-TD** |  | **NS-TD** | | **NF1-TD** |  | **NS-TD** | | **NF1-TD** |  | **NS-TD** | | **NF1-TD** |
| **CC body** | | | | | | | | | | |  | | | |  | | | |
| ***central*** | | **.004** | | **.002** | | **.037** | .071 |  | |  | **<.001** | **<.001** | | ***<*.001** | **<.001** | **<.001** | | **<.001** |
| ***parietal*** | | **<.001** | | **.008** | | **.001** | **.005** | **.010** | | **.008** | **<.001** | **<.001** | | ***<*.001** | **<.001** | **<.001** | | **<.001** |
| **prefrontal** | | **.017** | | .167 | | **.003** | .066 |  | |  | **<.001** | **.002** | | ***<*.001** | **<.001** | **<.001** | | **<.001** |
| **premotor** | | **.006** | | .081 | | **.002** | **.002** | **.001** | | **.030** | **<.001** | **<.001** | | ***<*.001** | **<.001** | **<.001** | | **<.001** |
| ***temporal*** | | **<.001** | | ***<*.001** | | **.014** | **<.001** | **.003** | | **.002** | **<.001** | **<.001** | | ***<*.001** | **<.001** | **<.001** | | **<.001** |
| ***CC genu*** | | **.003** | | **.047** | | **<.001** | .350 | 0.323 | | 0.333 | **<.001** | **.002** | | ***<*.001** | **<.001** | **.003** | | **<.001** |
| CC rostrum | | .294 | |  | |  | .175 | 0.316 | | 0.122 | .060 |  | |  | **<.001** | **.013** | | **.013** |
| ***CC splenium*** | | **.002** | | **.002** | | **.004** | .869 | 0.967 | | 0.642 | **<.001** | **<.001** | | **<.001** | **<.001** | **<.001** | | **<.001** |
| ***L ATR*** | | **.001** | | ***<*.001** | | **.002** | **<.001** | .074 | | **<.001** | **<.001** | ***<*.001** | | **<.001** | **<.001** | **<.001** | | **<.001** |
| ***R ATR*** | | **.001** | | **<.001** | | **<.001** | **.004** | .068 | | **.002** | **<.001** | ***<*.001** | | **<.001** | **<.001** | **<.001** | | **<.001** |
| **L AF** | | **.001** | | ***<*.001** | | .455 | **<.001** | **.012** | | **<.001** | **<.001** | **<.001** | | **<.001** | **<.001** | **<.001** | | **<.001** |
| **R AF** | | **.001** | | ***<*.001** | | .694 | **<.001** | **.015** | | **<.001** | **<.001** | **<.001** | | **<.001** | **<.001** | **<.001** | | **<.001** |
| L FAT | | .183 | |  | |  | **<.001** | **<.001** | | **<.001** | **<.001** | **<.001** | | **.003** | **<.001** | **<.001** | | **<.001** |
| **R FAT** | | **.009** | | **.002** | | .218 | **<.001** | **.004** | | **<.001** | **<.001** | **<.001** | | ***<*.001** | **<.001** | **<.001** | | **<.001** |
| ***L ILF*** | | **.001** | | ***<*.001** | | **.001** | **<.001** | **.003** | | **<.001** | **<.001** | **<.001** | | ***<*.001** | **<.001** | **<.001** | | **<.001** |
| ***R ILF*** | | **.001** | | ***<*.001** | | **.003** | **<.001** | **.007** | | **.001** | **<.001** | **<.001** | | ***<*.001** | **<.001** | **<.001** | | **<.001** |
| ***L MLF*** | | **.001** | | ***<*.001** | | **.002** | **.002** | .128 | | **.005** | **<.001** | **<.001** | | ***<*.001** | **<.001** | **<.001** | | **<.001** |
| **R MLF** | | **.001** | | ***<*.001** | | .1 | **<.001** | **.006** | | **<.001** | **<.001** | **<.001** | | **<.001** | **<.001** | **<.001** | | **<.001** |
| ***L SLF I*** | | **.013** | | **.012** | | **.045** | **<.001** | **.001** | | **<.001** | **<.001** | **<.001** | | ***<*.001** | **<.001** | **<.001** | | **<.001** |
| **R SLF I** | | **.003** | | **.001** | | .067 | **<.001** | **.005** | | **<.001** | **<.001** | **<.001** | | ***<*.001** | **<.001** | **<.001** | | **<.001** |
| **L SLF II** | | **.001** | | **<.001** | | .369 | **<.001** | **.025** | | **<.001** | **<.001** | **<.001** | | ***<*.001** | **<.001** | **<.001** | | **<.001** |
| **R SLF II** | | **.002** | | **<.001** | | .677 | **<.001** | **.016** | | **<.001** | **<.001** | **<.001** | | **.001** | **<.001** | **<.001** | | **<.001** |
| **L SLF III** | | **.001** | | **<.001** | | .228 | **<.001** | **.001** | | **<.001** | **<.001** | **<.001** | | ***<*.001** | **<.001** | **<.001** | | **<.001** |
| **R SLF III** | | **.001** | | **<.001** | | .269 | **<.001** | **.003** | | **<.001** | **<.001** | **<.001** | | ***<*.001** | **<.001** | **<.001** | | **<.001** |
| **L UF** | | **.003** | | **.002** | | .087 | **.002** | .099 | | **.011** | **<.001** | **<.001** | | **.004** | **<.001** | **<.001** | | **<.001** |
| **R UF** | | **.004** | | **.006** | | .337 | **.003** | .067 | | **.006** | **<.001** | **<.001** | | .027 | **<.001** | **<.001** | | **.003** |

F-statistic from ANCOVA tests (*) FDR-adjusted p-values significant at an alpha level of 0.05 or less are bolded. Significant p-values from post hoc t-tests (NF1-TD; NS-TD) are bolded. The names of tracts affected in both groups relative to TD controls are italicized. AF: Arcuate Fasciculus; ATR: Anterior Thalamic Radiation; CC: Corpus Callosum; df: degrees of freedom; FAT: Frontal Aslant Tract; ILF: Inferior Longitudinal Fasciculus; MLF: Middle Longitudinal Fasciculus; NF1: Neurofibromatosis type 1; NS: Noonan syndrome; R: Right; SLF: Superior Longitudinal Fasciculus; TD: typically developing; UF: Uncinate Fasciculus

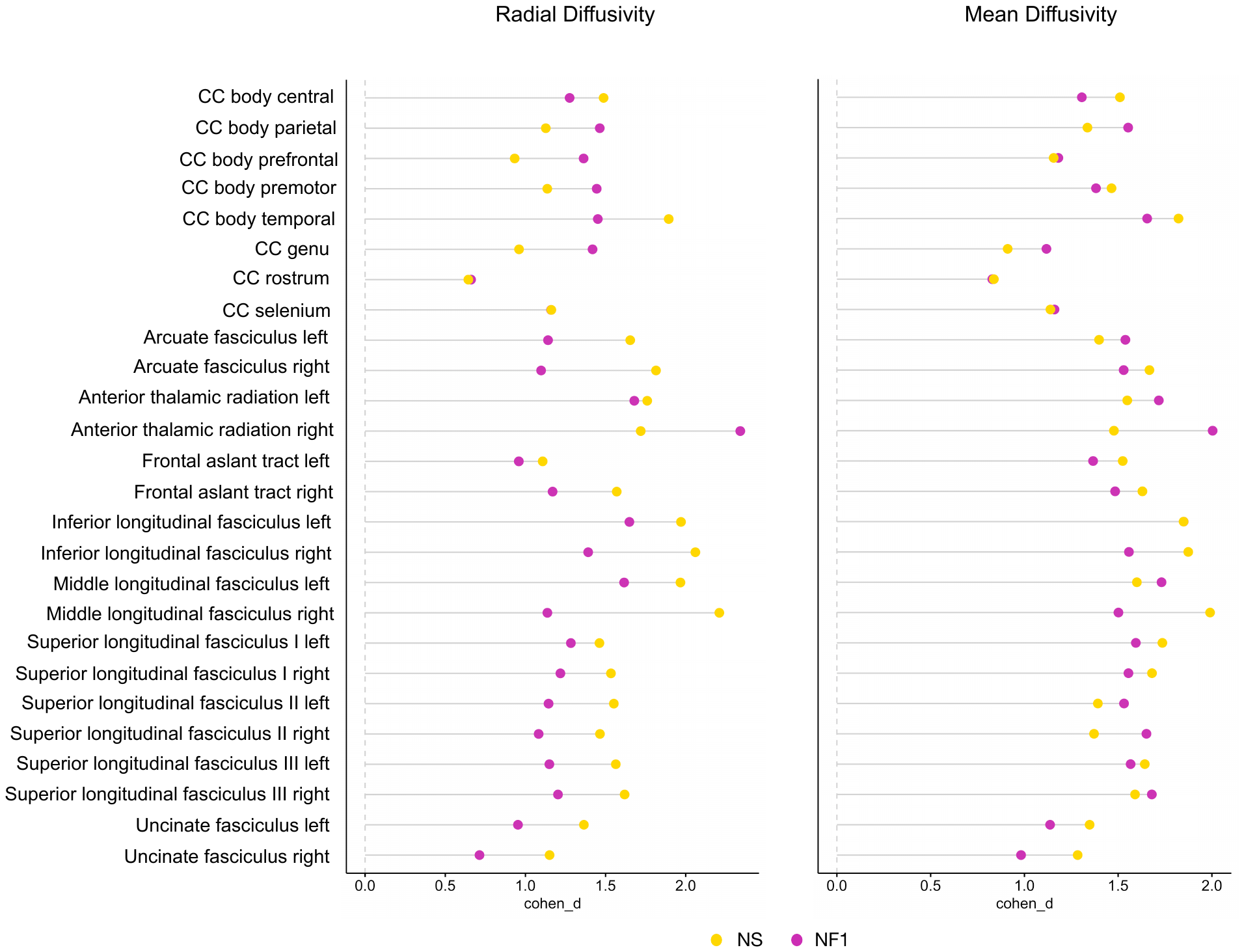

**Figure S1. Noonan syndrome (NS) and Neurofibromatosis type-1 (NF1) have similarly pervasive effects on neural white matter as indicated by radial (RD) and mean (MD) diffusivity.**

Effect sizes (cohen’s *d*) of mean differences between clinical groups and TD in RD (left) and MD (right) from select commissural, projection, and association tracts. NS’s cohen’s *d* values are depicted as yellow circles and NF1’s cohen’s *d* values are depicted as magenta circles.

### Supplementary references

1.	Noll DC, Nishimura DG, Macovski A. Homodyne detection in magnetic resonance imaging. *IEEE Trans Med Imaging*. 1991;10(2):154-163.

2.	Andersson JLR, Skare S, Ashburner J. How to correct susceptibility distortions in spin-echo echo-planar images: application to diffusion tensor imaging. *Neuroimage*. 2003;20(2):870-888.

3.	Andersson JLR, Graham MS, Zsoldos E, Sotiropoulos SN. Incorporating outlier detection and replacement into a non-parametric framework for movement and distortion correction of diffusion MR images. *Neuroimage*. 2016;141:556-572.

4.	Andersson JLR, Sotiropoulos SN. An integrated approach to correction for off-resonance effects and subject movement in diffusion MR imaging. *Neuroimage*. 2016;125:1063-1078.

5.	Andersson JLR, Graham MS, Drobnjak I, Zhang H, Filippini N, Bastiani M. Towards a comprehensive framework for movement and distortion correction of diffusion MR images: Within volume movement. *Neuroimage*. 2017;152:450-466.

6.	Andersson JLR, Graham MS, Drobnjak I, Zhang H, Campbell J. Susceptibility-induced distortion that varies due to motion: Correction in diffusion MR without acquiring additional data. *Neuroimage*. 2018;171:277-295.

7.	Jbabdi S, Sotiropoulos SN, Savio AM, Graña M, Behrens TEJ. Model-based analysis of multishell diffusion MR data for tractography: How to get over fitting problems. *Magn Reson Med*. 2012;68(6):1846-1855.

8.	Jenkinson M, Beckmann CF, Behrens TEJ, Woolrich MW, Smith SM. FSL. *Neuroimage*. 2012;62(2):782-790.

9.	*Non-Linear Registration, Aka Spatial Normalisation. FMRIB Technical Report TR07JA2*.; 2010. https://fsl.fmrib.ox.ac.uk/fsl/fslwiki/FNIRT

10.	Yendiki A, Panneck P, Srinivasan P, et al. Automated probabilistic reconstruction of white-matter pathways in health and disease using an atlas of the underlying anatomy. *Front Neuroinform*. 2011;5:23.

11.	Maffei C, Lee C, Planich M, et al. Using diffusion MRI data acquired with ultra-high gradient strength to improve tractography in routine-quality data. *Neuroimage*. 2021;245:118706.

12.	Maffei C, Gilmore N, Snider SB, et al. Automated detection of axonal damage along white matter tracts in acute severe traumatic brain injury. *NeuroImage Clin*. 2023;37(103294):103294.

13.	Yendiki A, Koldewyn K, Kakunoori S, Kanwisher N, Fischl B. Spurious group differences due to head motion in a diffusion MRI study. *Neuroimage*. 2014;88:79-90.
